# The association between reserve, cognitive ability and performance-related brain activity during episodic encoding and retrieval across the adult lifespan

**DOI:** 10.1101/535674

**Authors:** Abdelhalim Elshiekh, Sivaniya Subramaniapillai, Sricharana Rajagopal, Stamatoula. Pasvanis, Elizabeth Ankudowich, M. Natasha Rajah

## Abstract

Remembering associations between encoded items and their contextual setting is a feature of episodic memory. Although this ability generally deteriorates with age, there is substantial variability in how older individuals perform on episodic memory tasks. This variability may stem from genetic and/or environmental factors related to reserve, allowing some individuals to compensate for age-related decline through differential recruitment of brain regions. In this fMRI study spanning a large adult lifespan sample (N=154), we used multivariate Behaviour Partial Least Squares (B-PLS) analysis to examine how age, retrieval accuracy, and a proxy measure of reserve (i.e., a composite of years of education and premorbid I.Q.), impacted brain activity patterns during spatial and temporal context encoding and retrieval. We also conducted a secondary B-PLS to explore whether higher cognitive ability was associated with additional compensatory patterns capturing other aspects of reserve not accounted for by our composite measure of education and I.Q. Our results showed that higher reserve did not moderate the effect of age on cognitive ability or context memory, but higher cognitive ability was associated with better context memory performance and with task-specific compensatory responses in ventral visual, temporal, and fronto-parietal regions in advanced age. Additionally, higher reserve was associated with task-general responses in superior temporal, occipital, and inferior frontal regions. These findings suggest that task-specific compensatory responses in the aging brain are modulated by differences in general cognitive ability, which may be related to accumulated reserve, but not by proxy measures of reserve based on premorbid I.Q. and years of education.

## 1. Introduction

In everyday life we are commonly faced with instances where we need to remember past events that occurred at a specific time and place; such as, running into an acquaintance at the grocery store and trying to remember where we had initially met them. This type of long term memory for personally experienced events is referred to as episodic memory (Tulving, 2002). Episodic memory contains information about the content of past events, or item memory, and the surrounding details, such as the *when* and *where* of an event; commonly referred to as context/source memory (Johnson, Hashtroudi, & Lindsay, 1993; Tulving, 2002). Functional neuroimaging studies examining the neural underpinnings of successful context memory using face stimuli (i.e., face-location and/or temporal recency decisions) in younger adults have demonstrated that successful context memory relies on the activation of brain regions related to face processing (i.e., posterior ventral visual regions), prefrontal cortex (PFC), the hippocampus and surrounding MTL cortices, and parietal cortical regions (DuBrow & Davachi, 2014; Rajah, Ames, & D’Esposito, 2008; Rajah, Languay, & Valiquette, 2010; Sweegers & Talamini, 2014; Takashima et al., 2009, 2007).

In general, healthy aging is associated with a general decline in cognitive function (Park et al., 2002, 1996; Schaie, 2005). With respect to episodic memory, older adults show greater declines in context memory, compared to item memory (Spencer & Raz, 1995). However, most cognitive aging studies of context memory have focused on mean changes in memory performance with age, and thus assume that older adults (OA) are a homogenous group (Anderson et al., 2008; Cansino et al., 2013; Hashtroudi, Johnson, & Chrosniak, 1989; McIntyre & Craik, 1987; Wegesin, Jacobs, Zubin, Ventura, & Stern, 2000). Yet, there is significant variability in age-related context memory decline, and some OA perform comparably to young adults (YA) on some memory tasks (Christensen et al., 1999; Lindenberger & Ghisletta, 2009; Morse, 1993; Nilsson et al., 1997; Spreng, Wojtowicz, & Grady, 2010; Wilson et al., 2002; see Tucker-Drob & Salthouse, 2013 for a review). Indeed, several neuroimaging studies have examined the functional neural correlates that differentiate OA who perform similarly to YA on episodic memory tasks, compared to OA who exhibit the more traditional age-related episodic memory decline (Cabeza, Anderson, Locantore, & McIntosh, 2002; Duarte, Henson, & Graham, 2008; Meusel, Grady, Ebert, & Anderson, 2017). For example, Cabeza et al. (2002) compared brain activity in YA and OA, using Positron Emission Tomography (PET) during paired associate cued recall and context memory tasks. The OA were divided into high- and low-performance groups, based on their scores on a composite measure of frontal lobe function (Glisky, Polster, & Routhieaux, 1995). High-performing older adults (High-OA) showed matched performance to YA on both memory tasks, while low-performing older adults (Low-OA) were impaired on both tasks compared to YA. The PET results indicated that Low-OA showed greater right lateralized PFC activity during context memory, compared to the recall task. Moreover, this pattern of brain activity was similar to that observed in YA. In contrast, High-OA showed bilateral PFC engagement. This pattern of bilateral PFC activity in the High-OA group was interpreted as reflecting successful functional compensation in the aging brain, for underlying age-related deficits in PFC structural integrity (Cabeza & Dennis, 2013), or processing deficits in posterior cortical regions (Davis, Dennis, Daselaar, Fleck, & Cabeza, 2008; Grady et al., 1994). Therefore, variability in memory performance in advanced age may in part be related to variability in OA’s ability to engage neural compensatory mechanisms in response to structural/functional decline in other brain regions or networks.

The ability of some OA to perform as well as YA on memory tasks may also be explained by individual differences in *reserve* (Stern, 2002, 2012). The operational definitions of reserve, and closely related concepts, are still being developed and debated in the field today (Cabeza et al., 2018; Stern, Arenaza-Urquijo, et al., 2018). In the current manuscript, we define reserve as the accrual of neural resources over one’s lifetime, due to genetics and life experiences (environment), which help offset/attenuate the negative effects of age-related neural decline, and/or neuropathology, on cognitive function in later life (Cabeza et al., 20018; Stern et al., 2018). There are various potential neural mechanisms that may support reserve, as such, it is arguable that “reserve” itself is not a single entity and may not be directly measurable (Cabeza et al., 2018). However, there is evidence that some lifestyle and biological factors help support cognition in late life, i.e. education, intelligence, participation in leisure activities, and occupational complexity. These variables are typically used as indirect proxy measures for reserve, and cross-sectional studies indicate that older adults who have high levels of these proxy measures of reserve, exhibit better episodic memory performance (Angel, Fay, Bouazzaoui, Baudouin, & Isingrini, 2010; Lachman, Agrigoroaei, Murphy, & Tun, 2010). It has been hypothesized that having higher levels of these proxy measures of reserve may result in having greater neural capacity and increased availability and accessibility of neurocognitive strategies to perform various behavioural tasks; greater flexibility in the engagement of different neurocognitive strategies; and, greater neural efficiency in the utilization of these neurocognitive strategies and related brain regions and networks (Barulli & Stern, 2013). In other words, in relation to fMRI measurements of brain activity, an individual with higher proxy measures of reserve may be able to show less recruitment of task-related regions to perform a given task without compromising performance (i.e., efficiency); be able to maximize recruitment of task-related regions under increasing demands (i.e., capacity); or be able to utilize alternate networks to maintain or improve performance (i.e., flexibility). In that regard, functional compensation may be thought of as enhanced neural capacity and flexibility and may be supported by reserve in old age.

Consistent with this hypothesis, previous studies have shown differential recruitment of brain networks in older adults with high, compared to lower levels of proxy measures of reserve (Stern, 2012). For example, in an fMRI study, Springer et al. (2005) used multivariate partial least squares (PLS) analysis (McIntosh et al., 2004) to investigate the relationship between brain activity and years of education during episodic encoding and recognition in a group of YA and OA. In YA, they found that education and memory performance were positively correlated with activity in medial temporal, ventral visual and parietal cortices, and negatively correlated with activity in prefrontal cortex (PFC). In OA, higher education was related to increased activity in bilateral PFC and right parietal cortex; however, this pattern of brain activity was not directly correlated with better memory performance in OA. One possible interpretation of these findings is that YA with higher reserve exhibit greater neural efficiency in PFC activation, and greater neural capacity in medial temporal, ventral visual and parietal cortices. In contrast, older age is associated with reduced neural efficiency in PFC function and reduced capacity in parietal function.

More recently, Stern et al. (2018) examined blocked and event-related task fMRI data from a variety of cognitive domains in 58 young adults (aged 18-31) and 91 older adults aged (51-71) and identified a general pattern of brain activation that varied with intelligence quotient (I.Q.), as measured by the North American Reading Test (NART; (Nelson & Wilison, 1991)) (Stern et al., 2018). They found that increased activity in cerebellum, medial PFC, and bilateral superior frontal gyrus across all tasks was associated with having higher IQ. They also found that higher IQ was related to decreased activity in bilateral middle and inferior prefrontal cortex PFC and bilateral inferior parietal cortex. However, the correlation between IQ-related brain activity and task performance and age was not investigated. This suggests that higher IQ is associated with greater efficiency in lateral PFC and inferior parietal cortex, and increased capacity in cerebellum and medial superior PFC.

Taken together, the results from Cabeza et al (2002), Springer et al (2005) and Stern et al (2018) suggest that higher cognitive ability and/or reserve may be related to greater efficiency in lateral PFC function and increased capacity in medial superior PFC and posterior cortical regions, including medial temporal and ventral visual cortices (Stern et al 2018; Springer et al 2005). Moreover, the results from Cabeza et al (2002) and Springer et al (2005) indicate that age-related memory decline may be related to increased neural inefficiency in lateral PFC with age; and, that older adults with greater cognitive ability (and possibly reserve) may have greater neurocognitive flexibility that promotes the activation of additional PFC regions to support episodic memory in late life. These findings raise the question: are reserve-related increases in neural efficiency, neural capacity and flexibility observed in distinct brain regions through out the adult lifespan? Or, is the association between reserve, and observed decreases (neural efficiency) and increases (capacity and/or flexibility) in task-related brain activity differ across the adult lifespan and differ depending on task demands/difficulty (Stern et al., 2018). Finally, it remains unclear if reserve-related modulation in brain activity directly supports episodic memory performance, and if such an effect is stable across the adult lifespan. In other words, it remains unclear if there is a positive association between proxy measures of reserve, level of cognitive function, and increases or decreases in brain activity to support performance on a variety of episodic memory tasks, at varying levels of difficulty, across the adult lifespan. In the current study we test this hypothesis.

In this study, 154 adults between the ages of 19-76 years underwent neuropsychological testing and fMRI scanning during easy and difficult versions of left/right face-location spatial context memory tasks and least/most recent face temporal context memory task. FMRI scans were obtained during both encoding and retrieval. Initial analyses that explored age and performance-related patterns of brain activity in a subset of this dataset (N = 128) have been previously published (Ankudowich, Pasvanis, & Rajah, 2016, 2017). Here, we tested 26 more adults in this experimental paradigm. We calculated a composite measure of cognitive ability, based on neuropsychological assessments; and a proxy measure of reserve, based on years of education and performance on the NART, a measure of crystallized I.Q. (Nelson & Wilison, 1991). We then tested the hypothesis that reserve moderated the effect of age on our composite measure of cognitive ability and measures of episodic memory function obtained from task fMRI and used multivariate behaviour partial least squares (B-PLS) to examine how age, reserve, and memory performance related to brain activity at encoding and retrieval across the adult lifespan. We predicted that during easy tasks we would observe that greater reserve would be related to decreased activity (greater neural efficiency) in lateral PFC and parietal regions (Stern et al., 2018), and increased activity (greater neural capacity / flexibility) in additional PFC regions, medial temporal and ventral visual regions, across the adult lifespan; and this in turn would related to better episodic memory ability (Ankudowich et al., 2017; Cabeza et al., 2002; Springer et al., 2005). In contrast, we hypothesized that during difficult memory tasks we would observe increased activity in lateral PFC and parietal cortices (Stern et al., 2018), which may support better memory performance on these more challenging tasks across the adult lifespan.

## 2. Methods

### 2.1 Participants

One hundred and fifty-four healthy adults (age range 19-76 yrs, mean age = 48.08 yrs; 109 females; mean years of formal education [EDU] = 15.66 yrs) participated in this study. Of the 154 participants tested, 42 were young (age range 19-35 yrs, mean age = 25.81 yrs; 28 females; EDU = 16.21 yrs), 68 were middle-aged (age range 40-58 yrs, mean age = 50.00 yrs; 51 females; EDU = 15.35 yrs), and 44 were old (age range 60-76 yrs, mean age = 66.39 yrs; 30 females; EDU = 15.61 yrs). The age groups did not differ in level of education. All participants were right-handed as assessed by the Edinburgh Inventory for Handedness (Oldfield, 1971), had no history of neurological or psychological illness, and had no family history of Alzheimer’s disease.

Participation involved two sessions conducted on two separate days. During the first session, participants completed a battery of neuropsychological measures assessing their eligibility. The measures consisted of the Folstein Mini Mental State Examination (MMSE), exclusion cut-off < 27, (Folstein, Folstein, & McHugh, 1975); the Beck Depression Inventory (BDI), exclusion cut-off > 15, (Beck, Ward, Mendelson, Mock, & Erbaugh, 1961), the NART used as a proxy for premorbid crystallized intelligence, inclusion cut-off ≤ 2.5 SD, (Nelson & Wilison, 1991), and a practice run of the context memory tasks in a mock MRI scanner. The California Verbal Learning Test (CVLT) Delay Free Recall (DFR), CVLT Delay Cued Recall (DCR), and CVLT Delay Recognition (DRG) was also administered to assess long term memory recall (Delis, Kramer, Kaplan, & Thompkins, 1987). To assess executive function, the Wisconsin Card Sorting Test (WCST; Heaton, Chelune, Talley, Kay, & Curtiss, 1993) and the Delis-Kaplan Executive Function System (D-KEFS; Delis, Kaplan, & Kramer, 2001) verbal fluency test; Letter Fluency (LF), Category Fluency (CF), and Category Switching (CS), were also administered. Additional medical exclusion criteria included: having a history of diabetes, untreated cataracts and glaucoma, smoking > 40 cigarettes a day, and a current diagnosis of high cholesterol and/or high blood pressure that has been left untreated in the past six months. Individuals who met the neuropsychological inclusion criteria and performed above chance on the mock-MRI scanner trials, were invited to participate in a second fMRI testing session. All participants were paid and provided their informed consent to participate in the study. The ethics board of the Faculty of Medicine at McGill University approved the study protocol.

### 2.2 Behavioural methods

Details concerning the methods and stimuli pertaining to the fMRI task have previously been outlined in Kwon et al. (2016). In brief, a mixed rapid event-related fMRI design was implemented in which participants were scanned while encoding and retrieving the spatial context (whether a face had appeared on the left or the right side of the screen during encoding) or temporal context (whether a face had appeared most or least recently at encoding) of face stimuli. Participants completed 12 experimental runs of easy and difficult versions of the spatial and temporal tasks. Each run consisted of one spatial easy (SE) and one temporal easy (TE) context memory task, in addition to either a spatial hard (SH), or temporal hard (TH) task. During easy tasks, participants encoded 6 face stimuli and during hard tasks participants encoded 12 face stimuli.

The task stimuli set has been used in previous studies (Rajah et al., 2008, 2010), and consisted of black and white photographs of age variant faces. All face stimuli were cropped at the neck and were rated for pleasantness by two independent raters. The age and sex of the faces were balanced across experimental conditions and were each presented only once at encoding without replacement. Faces shown at encoding were subsequently tested at retrieval.

#### 2.2.1 Encoding phase

At the start of each encoding phase, participants were cued (9s) to memorize either the spatial location or temporal order of the ensuing faces, then either six (i.e., easy) or 12 (i.e., hard) faces were serially presented to the left or right side of a centrally presented fixation cross. Each stimulus was presented for 2s followed by a variable ITI (2.2-8.8s). Participants were also asked to rate the pleasantness of each face as pleasant or neutral during encoding. This was done to ensure subjects were on task and encoded the faces. In total, participants performed 12 SE, 12 TE, 6 SH, and 6 TH tasks, yielding a total of 72 encoding events per task-type (i.e., 288 total encoding events). Following the encoding phase of each run, participants performed an alphabetization distraction task (60s) where they were asked to select the word that comes first in the alphabet. The distraction task served to minimize working-memory related rehearsal of encoded information.

#### 2.2.2 Retrieval phase

After the distraction task, participants were cued (9s) that the retrieval phase (spatial or temporal) was about to begin. Depending on the retrieval task cued, participants were presented with two previously encoded faces above and below a central fixation cross and were either asked which face was originally presented to the left (or right) side of the screen during encoding (spatial context retrieval), or was originally seen least/most recently (temporal context retrieval). Easy retrieval tasks consisted of three retrieval pairs and hard retrieval tasks consisted of six retrieval pairs, for a total of 36 retrieval events per task type. Each retrieval pair was presented for 6s followed by a variable ITI (2.2-8.8s).

### 2.3 Behavioural data analysis

SPSS version 24 (IBM Corp., 2016) was used to conduct repeated-measures mixed effects ANOVAs on retrieval accuracy (% correct) and reaction time (msec) with group (3: young, middle-aged, older adults) as a between-subjects factor, and task (2: spatial, temporal) and difficulty (2: easy, hard) as within-subject factors, to determine significant group, task and difficulty main effects, and interactions (significance threshold p < 0.05). Post-hoc tests were conducted as needed to clarify significant effects and interactions.

#### Calculation of composite measure of cognitive ability and proxy measure of reserve

The composite measure of cognitive ability was based on neuropsychological test data. Every participant’s raw score on the CVLT-DFR, DCR, DRG; D-KEFS-LF, CF, CS; WCST categories completed, and correct responses (%) were standardized by z-score transformation based on the full sample. An average standardized score was then computed for every participant based on the z-scores of the eight items mentioned above to yield a composite cognitive ability score. These items were selected because we were interested in creating an overall cognitive ability measure that provides a proxy for memory ability (CVLT; Delis et al., 1987) and executive function (D-KEFS, and WCST; Delis et al., 2001; Heaton et al., 1993).

We also calculated a proxy measure of reserve for each participant by calculating the mean value of z-scored education in years and z-scored estimated premorbid crystallized I.Q. based on the NART.

#### Regression analysis

To test the hypothesis that reserve moderated the effect of age on our composite measure of cognitive ability and the task fMRI accuracy and RT measures, we used linear regression implemented in R (R Core Team, 2014) to test the following models: DV ∼ β + Age + Reserve + Age*Reserve + ε, in which DV = cognitive ability; mean accuracy on SE, SH, TE and TH tasks; and mean RT for SE, SH, TE and TH tasks. Significance assessed at p < 0.05 corrected for multiple comparisons.

Given that our proxy measure of reserve may not fully capture the factors that support successful cognitive aging, we also conducted exploratory regression analyses to test the hypothesis that our composite measure of cognitive ability moderated the effect of age on task fMRI accuracy and RT measures using the model: DV ∼ β + Age + Cognitive Ability + Age*Cognitive Ability + ε, in which DV = mean accuracy on SE, SH, TE and TH tasks; and mean RT for SE, SH, TE and TH tasks. Significance was assessed at p < 0.05 corrected for multiple comparisons.

### 2.4 MRI methods

Structural and functional MRI scans were collected on a 3T Siemens Trio scanner at the Douglas Institute Brain Imaging Centre. Participants wore a standard 12-channel head coil while lying in supine position. T1-weighted anatomical images were acquired at the beginning of the fMRI testing session using a 3D gradient echo MPRAGE sequence (TR = 2300 ms, TE = 2.98 ms, flip angle = 9°, 176 1 mm sagittal slices, 1 × 1 × 1 mm voxels, FOV = 256 mm^2^). FMRI BOLD images were acquired using a single shot T2*-weighted gradient echo planar imaging (EPI) pulse sequence (TR = 2000 ms, TE = 30 ms, FOV = 256 mm^2^, matrix size = 64 × 64, in plane resolution 4 × 4 mm, 32 oblique 4 mm slices with no slice gap) during the context memory task. Jitter was added to the event-related acquisitions by means of a mixed rapid event related design with variable ITI (as stated above).

Visual task stimuli were back projected onto a screen in the scanner bore and was made visible to participants lying in the scanner via a mirror mounted within the standard head coil. The stimuli were generated on a computer using E-Prime (Psychology Software Tools, Inc.; Pittsburgh, PA, USA) software. Participants requiring visual acuity correction wore corrective plastic lenses and a fiber optic 4-button response box was supplied to participants to make responses during the task.

#### 2.4.1 Pre-processing

Images were converted from DICOM to ANALYZE format using Statistical Parametric Mapping (SPM) version 8 software (http://www.fil.ion.ucl.ac.uk/spm) run with MATLAB (www.mathworks.com). SPM8 was used for pre-processing on a Linux platform. To ensure that all tissue had reached steady state magnetization, images acquired during the first 10s were discarded from analysis. The origin of functional images for each participant was reoriented to the anterior commissure of the T1-weighted structural image. Functional images were then realigned to the first BOLD image and corrected for movement using a 6 parameter rigid body spatial transform and a least squares approach. One participant had > 4mm movement and was excluded from further analysis. Functional images were then spatially normalized to the MNI EPI template (available in SPM) at 4 × 4 × 4 mm voxel resolution, and smoothed with an 8 mm full-width half maximum (FWHM) isotropic Gaussian kernel. ArtRepair toolbox for SPM8 was used to correct for bad slices prior to realignment and for volume artificats after normalization and smoothing (http://cibsr.stanford.edu/tools/human-brain-project/artrepair-softwarte.html).

#### 2.4.2 Multivariate PLS analysis

We conducted a Multivariate Behavioural PLS (B-PLS) analysis to identify how whole brain patterns of activity varied as a function of age, reserve, and/or task accuracy at encoding and retrieval. We selected B-PLS for our analyses due to its ability to identify spatially and temporally distributed voxel activation patterns that are differentially related to the experimental conditions and/or correlated with the behavioural vectors of choice (McIntosh, Chau, & Protzner, 2004). The first step in B-PLS was to represent the fMRI data for correctly encoded and retrieved events in an fMRI data matrix. To do this, the three-dimensional event-related fMRI data were converted to a two-dimensional data matrix by ‘flattening’ the temporal dimension (t), so that time series of each voxel (m) is stacked side-by-side across the columns of the data matrix (column dimension = m*t) (McIntosh et al., 2004). The rows of the 2D data matrix reflect the following experimental conditions nested within subjects: SE encoding, SH encoding, TE encoding, TH encoding, SE retrieval, SH retrieval, TE retrieval, TH retrieval. The columns of the fMRI data matrix reflect the event-related activity for each brain voxel, at each time point, for correctly encoded and retrieved events. For each event, activity was included for seven time-points/measurements, equivalent to 7 TRs (TR = 2 sec * 7 = 14 sec of activity per event), following the event onset. Thus, the first column of the data matrix reflected brain activity at event onset; the second column of the data matrix reflected activity at 2 sec following the event onset; the third column of the data matrix reflected activity at 4 sec following the event onset; and so forth. To control for low frequency signal drifts due to environmental and/or physiological noise (McIntosh et al., 2004), event-related activity was base-line corrected (zeroed) to the event onset. The event-related brain activity is then mean-centred within condition. As such, the data matrix reflected mean corrected percent change in brain activity from event onset for all conditions, stacked within subjects.

The fMRI data matrix was then cross-correlated with three behavioural vectors stacked in the same condition, nested within subject order: age, proxy measure of reserve (reserve), and mean retrieval accuracy per condition. The mean retrieval accuracy included in the analysis was orthogonalized to the age variable by obtaining its residual from a linear regression in which age was the predictor. This was done because age and raw accuracy were correlated. The resulting cross-correlation matrix was then submitted to singular value decomposition (SVD), which yielded mutually orthogonal latent variables (LVs). Each LV consists of: i) a singular value, reflecting the amount of covariance explained by the LV; ii) a correlation profile, which reflects how the three behavioural vectors correlate with a pattern of whole-brain activity identified in the singular image (described next); iii) a singular image, which depicts a pattern of brain saliences, reflecting numerical weights assigned to each voxel at each TR/time lag included in the data matrix. These brain saliences represent a pattern of whole-brain activity that is symmetrically related to the correlation profiles for each of the three behavioural vectors. Brain regions with positive saliences are positively related to the correlation profile (with 95% confidence intervals), while those with negative saliences are negatively related to the correlation profile (with 95% confidence intervals). Since each LV reflects a symmetrical pairing of correlation profiles with a pattern of whole-brain activity, the inverse can also be implied; positive values in the correlation profile indicate a negative correlation with negative salience brain regions, and negative values indicate a positive correlation with negative salience brain regions.

Significance of LVs was assessed through 1000 permutations for each B-PLS analysis. The permutation test involved sampling without replacement to reassign links between subjects’ behavioural vector measures and event/condition within subject. For each permuted iteration a PLS was recalculated, and the probability that the permuted singular values exceeded the observed singular value for the original LV was used to assess significance at p < 0.05 (McIntosh et al., 2004). To identify stable voxels that consistently contributed to the correlation profile within each LV, the standard errors of the voxel saliences for each LV were estimated via 500 bootstraps, sampling subjects with replacement while maintaining the order of event types for all subjects. For each voxel, a value similar to a z-score known as the bootstrap ratio (BSR) was computed, reflecting the ratio of the original voxel salience to the estimated standard error for that voxel. Voxels with BSR values of ± 3.28 (equivalent to p < 0.001) and a minimum spatial extent = 10 contiguous voxels, were retained and highlighted in the singular image. BSR values reflect the stability of voxel saliences. A voxel salience whose value is dependent on the observations in the sample is less precise than one that remains stable regardless of the samples chosen (McIntosh & Lobaugh, 2004).

In order to determine at which time lags the correlation profile in a given LV was strongest, we computed temporal brain scores for each significant LV. Temporal brain scores reflect how strongly each participant’s data reflected the pattern of brain activity expressed in the singular image in relation to its paired correlation profile, at each time lag. Peak coordinates are only reported from time lags at which the correlation profile was maximal differentiated within the temporal window sampled (lags 2-5; 4-10s after event onset). These peak coordinates were converted to Talairach space using the icbm2tal transform (Lancaster et al., 2007) as implemented in GingerAle 2.3 (Eickhoff et al., 2009). Since our acquisition incompletely acquired the cerebellum, peak coordinates from this region are not reported. The Talairach and Tournoux atlas (Talairach & Tournoux, 1998) was used to identify the Brodmann area (BA) localizations of significant activations. To confirm our interpretations of the effects represented in each significant LV, we ran post-hoc GLM comparisons on the brain scores for the task conditions against our three behavioural vectors of interest.

## 3. Results

### 3.1 Behavioural results

Table 1 summarizes demographics, neuropsychological test data, and context memory performance for all groups. One-way ANOVAs indicated that there was a significant age-group difference in cognitive ability (F (2,151) = 8.197, p < 0.001, η2 = 0.098) due to the fact that younger and older adults scored higher than middle-aged adults on the cognitive ability composite measure. Similarly, both young and older adults had higher cognitive reserve compared to middle-aged adults (F (2,151) = 3.503, p < 0.033, η2 = 0.044).

**Table 1.** Demographics, neuropsychological test data, and context memory performance per age-group

| | Young | Middle-aged | Old | P-value | $\eta^2$ |
| --- | --- | --- | --- | --- | --- |
| **Sample size** | 42 | 68 | 44 |  |  |
| **Age (Yrs)** | 25.81 (0.54) | 50.00 (0.65) | 66.39 (0.56) |  |  |
| **Gender (n, [%] females)** | 28 [67%] | 51 [75%] | 30 [68%] |  |  |
| **Cognitive ability composite <sup>a</sup>** | 0.45 (0.12) | -0.29 (0.13) | 0.13 (0.13) | < .001* | .098 |
| **Reserve composite <sup>a</sup>** | 0.26 (0.13) | -0.20 (0.12) | 0.16 (0.15) | .033* | .044 |
| **EDU (Yrs)** | 16.21 (0.29) | 15.35 (0.25) | 15.61 (0.34) | .104 | - |
| **CVLT – DFR <sup>b</sup>** | 13.88 (0.27) | 12.75 (0.26) | 13.07 (0.33) | .021* | .050 |
| **CVLT – DCR** | 13.98 (0.27) | 13.13 (0.24) | 13.23 (0.30) | .072 | - |
| **CVLT – DRG** | 15.52 (0.10) | 15.13 (0.13) | 15.18 (0.13) | .084 | - |
| **WCST – categories completed <sup>c</sup>** | 8.52 (0.15) | 6.51 (0.31) | 6.32 (0.36) | < .001* | .154 |
| **WCST – % correct <sup>c</sup>** | 0.83 (0.07) | 0.73 (0.15) | 0.74 (0.17) | < .001* | .146 |
| **D-KEFS – LF** | 12.05 (0.46) | 11.64 (0.43) | 12.66 (0.52) | .302 | - |
| **D-KEFS – CF <sup>d</sup>** | 11.67 (0.50) | 10.94 (0.43) | 13.41 (0.41) | < .001* | .094 |
| **D-KEFS – CS** | 13.48 (0.47) | 13.38 (0.38) | 14.48 (0.36) | .131 | - |
| **Estimated I.Q. (NART)** | 119.62 (0.81) | 118.00 (0.73) | 120.35 (0.69) | .067 | - |
| **Accuracy (% correct)** |  |  |  |  |  |
| *Spatial easy retrieval* | 0.89 (0.01) | 0.86 (0.01) | 0.82 (0.01) |  |  |
| *Temporal easy retrieval* | 0.77 (0.20) | 0.70 (0.16) | 0.65 (0.01) |  |  |
| *Spatial hard retrieval* | 0.89 (0.15) | 0.81 (0.01) | 0.77 (0.02) |  |  |
| *Temporal hard retrieval* | 0.68 (0.02) | 0.59 (0.01) | 0.54 (0.01) |  |  |
| **Reaction time (msec)** |  |  |  |  |  |
| *Spatial easy retrieval* | 2198 (78.60) | 2501 (63.12) | 2781 (71.20) |  |  |
| *Temporal easy retrieval* | 2596 (84.51) | 2966 (59.94) | 3157 (88.97) |  |  |
| *Spatial hard retrieval* | 2303 (74.03) | 2628 (54.12) | 2837 (77.34) |  |  |
| *Temporal hard retrieval* | 2777 (95.58) | 3099 (75.70) | 3185 (93.30) |  |  |

Note: This table presents age-group means and standard errors between brackets for demographic, neuropsychological measures, and spatial and temporal context memory accuracy and reaction times. In addition, one-way ANOVA p-values and partial eta squared ( $\eta^2$ ) values for demographic and neuropsychological measures are listed. EDU = Years of Education; CVLT = California Verbal Learning Test; DFR = Delay Free Recall; DCR = Delay Cued Recall; DRG = Delay Recognition; WCST = Wisconsin Card Sorting test; D-KEFS = Delis-Kaplan Executive Function System; LF = Letter Fluency; CF = Category Fluency; CS = Category switching; NART = American National Adult Reading Test.
Tukey's HSD post-hoc between-group tests were conducted at $p = 0.05$ to clarify group differences and are summarized as follows: <sup>a</sup> young & old adults > middle-aged adults; <sup>b</sup> young adults > middle-aged adults; <sup>c</sup> young adults > middle-aged & old adults; <sup>d</sup> old adults > young & middle-aged adults.

On, the CVLT delay free recall, younger adults outperformed middle-aged adults (F (2,151) = 3.94, p = 0.021, η2 = 0.050), and completed more categories on the WCST (F (2,151) = 13.73, p < 0.001, η2 = 0.154) than both middle-aged and older adults. Younger adults were also more accurate on the WCST (F (2,151) = 12.96, p < 0.001, η2 = 0.146) than both middle-aged and older adults. On the D-KEFS category fluency, older adults outperformed both younger and middle-aged adults (F (2,151) = 7.79, p < 0.001, η2 = 0.094).

#### 3.1.1 Accuracy results

The group (3) × task (2) × difficulty (2) repeated-measures (RM) ANOVA on retrieval accuracy revealed a significant main effect of group (F (2, 151) = 22.19, p < 0.001, η2 = 0.227), task (F (1, 151) = 763.30, p < 0.001, η2 = 0.835), difficulty (F (1, 58) = 115.33, p < 0.001, η2 = 0.433), and a task × difficulty interaction (F (1, 151) = 34.76, p < 0.001, η2 = 0.187). Tukey’s HSD post-hoc test indicated that the significant main effect of group was due to younger adults outperforming both middle-aged and older adults, and middle aged adults outperforming older adults across conditions (ps < 0.001). Across groups, participants performed better on the spatial compared to the temporal task, and on easy vs. hard tasks. However, the significant task × difficulty interaction indicated that the difficulty manipulation impacted accuracy scores more on the temporal task (t (1,153) = 11.22, p < 0.001), compared to the spatial task (t (1,153) = 4.82, p < 0.001). We did not observe an overall age*difficulty interaction in the current study. This may be since the temporal context memory task was challenging to all age-groups. Exploratory repeated one-way ANOVAs examining age and difficulty effects within task-type verified this interpretation. For the temporal context memory tasks, we observed significant main effects of age-group (F (2, 151) = 22.66, p<0.001) and difficulty (F (1,151) =116.72, p<0.001), but no significant age*difficulty interaction. However, for the spatial context memory tasks we observed a significant age*difficulty interaction (F (2,151) = 4.08, p< 0.05); and, significant main effects of age-group (F (2,151) = 12.89, p<0.001) and difficulty (F (1, 151) = 20.12, p<0.001).

Within-age group correlations between the composite cognitive ability measure and retrieval accuracy for each task (i.e., SE, SH, TE, and TH) failed to reach significance threshold for any of the tasks per age group. Yet, cognitive ability was positively correlated with spatial easy (r (154) = 0.16, p = 0.045), spatial hard (r (154) = 0.17, p = 0.031), and temporal hard (r (154) = 0.23, p = 0.004) retrieval accuracy across participants. On the other hand, cognitive reserve was not significantly associated with task accuracy within- or across-age groups.

#### 3.1.2 Reaction time results

The group (3) × task (2) × difficulty (2) RM ANOVA on retrieval reaction time (RT) revealed a significant main effect of group (F (2, 151) = 12.93, p < 0.001, η2 = 0.146), task (F (1, 151) = 204.04, p < 0.05, η2 = 0.575), and difficulty (F (1, 58) = 31.19, p < 0.05, η2 = 0.171). Tukey’s HSD post-hoc test indicated that younger adults responded faster than both middle-aged and older adults across conditions (ps < 0.001). Across groups, participants were slower on the temporal compared to the spatial task, and on hard vs. easy tasks (ps < 0.001).

#### 3.1.3 Regression analyses results

The regression model for: Cognitive Ability ∼ β + Age + Reserve + Age*Reserve + ε was not significant (p > 0.05). Similarly, the regression models with task accuracy and reaction times for each of the task conditions (SE, SH, TE, TH), as dependant variables, and age, reserve, and age*reserve as predictors, did not yield any significant main effects of reserve, or age*reserve interactions. However, age was a significant predictor for all the models indicating that task performance decreased with advanced age. The lack of a main effect of reserve or age*reserve interaction indicates that our proxy measure of reserve did not modulate cognitive ability or memory performance.

In our current sample, age correlated negatively with cognitive ability (r = −0.20, p = 0.02), but not with reserve (p > 0.05). Therefore we suspected that our exploratory regression models with task accuracy and reaction times for each of the task conditions as dependant variables, and age, cognitive ability, and age*cognitive ability as predictors might suffer from multicollinearity. To assess for multicollinearity in our models, we computed a variance inflation factor (VIF), which revealed a score of 14.3 for cognitive ability and age. This score exceeds the acceptable threshold of 10 for multicollinearity in regressions models indicating redundancy between our predictor variables (Hair, Black, Babin, & Anderson, 2010; James, Witten, Hastie, & Tibshirar, 2017). To account for multicollinearity in our models, we orthogonalized the age variable to the cognitive ability variable by obtaining its residual from a linear regression in which cognitive ability was the predictor. We then ran our regression models with age-residuals, cognitive ability, and age-residuals*cognitive ability as predictor variables, and task accuracy and reactions times for each task condition as dependant variables.

We found a significant effect for the model predicting SE accuracy (F (3, 150) = 7.14, p < 0.001) with age (t = −4.10, p < 0.01) and cognitive ability (t = 2.28, p = 0.02) as significant predictors in the model. Similarly, the model for SH accuracy was significant (F (3, 150) = 12.16, p < 0.001) with age (t = −5.42, p < 0.01) and cognitive ability (t = 2.42, p = 0.02) as significant predictors. The model for TE accuracy was significant (F (3, 150) = 9.21, p < 0.001) with only age predicting accuracy (t = −5.11, p < 0.01), and the model for TH accuracy was significant (F (3, 150) = 19.31, p < 0.001) with both age (t = −6.85, p < 0.01) and cognitive ability (t = 3.45, p < 0.01) as significant predictors in the model. For reaction times, we found a significant effect for the model predicting SE reaction times (F (3, 150) = 14.08, p < 0.001) with both age (t = 5.79, p < 0.01) and cognitive ability (t = −3.07, p < 0.01) as significant predictors in the model. Likewise, the model for SH reaction times was significant (F (3, 150) = 12.57, p < 0.001) with both age (t = 5.54, p < 0.01) and cognitive ability (t = −2.67, p < 0.01) as significant predictors. The model for TE reaction times was significant (F (3, 150) = 10.3, p < 0.001) with only age as a significant predictor (t = 5.15, p < 0.01) and cognitive ability approaching significance (t = −1.80, p = 0.07). The model for TH reaction times was also significant (F (3, 150) = 4.47, p < 0.001), with age as the only significant predictor (t = 3.27, p < 0.01). Therefore, with respect to cognitive ability, our results indicated that higher cognitive ability was associated with more accurate responses for all task conditions (except for TE), and quicker reaction times during SE and SH task performance.

### 3.2 fMRI results

#### 3.2.1 B-PLS: Age, Reserve and Accuracy

The B-PLS analysis revealed five significant LVs linking whole-brain patterns of activity to the behavioural vectors of age, reserve, and task accuracy (residualized by age). LV1 accounted for 19.76% of the total cross-block covariance (p < 0.001). Only negative salience brain regions from this LV survived our spatial threshold cut-off of 10 contiguous voxels (p < 0.001), and the local maxima of those negative saliences are presented in Table 2. Figure 1a shows the PLS correlation profile separated by task (SE, SH, TE, and TH), and the corresponding singular image presented in Figure 1b demarcates the stable negative salience regions (cool coloured regions). The PLS correlation profile indicates that this LV was mostly related to easy events across both tasks (spatial and temporal). Specifically, activity in negative salience brain regions increased with age during easy encoding events (SE, and TE). Activity in those regions was also correlated positively with subsequent accuracy, (but not with reserve) for the same easy spatial and temporal encoding events. Interestingly, activity in negative salience regions was also positively correlated with accuracy at easy spatial and temporal retrieval events. In other words, LV1 primarily identified negative salience brain regions in which event-related activity increased with age, and subsequent retrieval accuracy during easy encoding events, and increased with retrieval accuracy during easy retrieval events.

**Figure 1.**
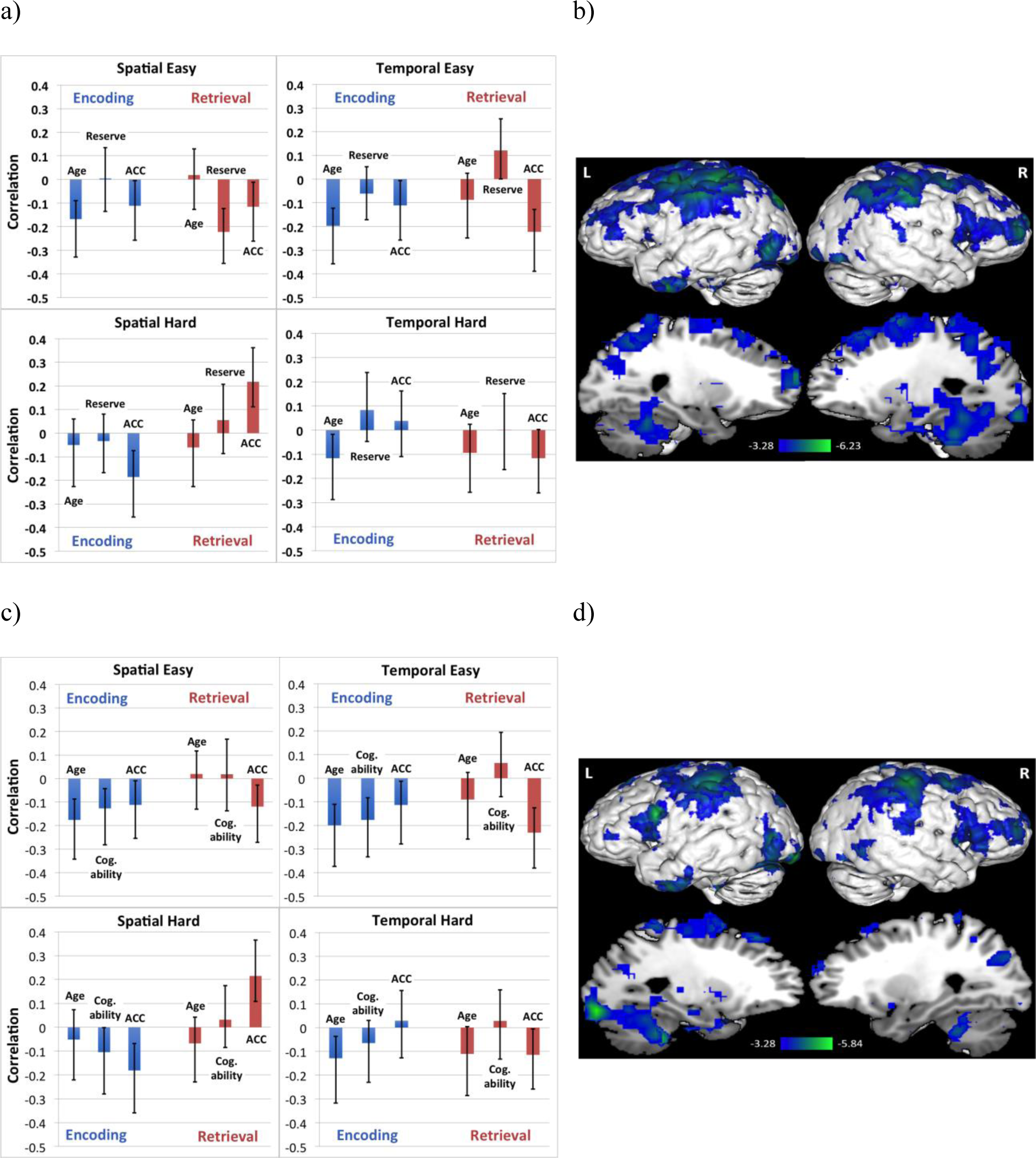
Brain-behaviour correlation profiles and corresponding singular images for LV1 in both B-PLS analysis. a) LV1 brain-behaviour correlation profile for reserve B-PLS separated by task. The correlation profile indicated that activity in negative salience brain regions correlated positively with age, and accuracy (ACC) during easy encoding events, and correlated positively with ACC during easy retrieval events. c) LV1 brain-behaviour correlation profile for cognitive ability B-PLS. The correlation profile indicated that activity in negative salience brain regions correlated positively with age, cognitive ability (Cog. ability), and ACC during easy encoding events, and correlated positively with ACC during easy retrieval events. Error bars represent 95% confidence intervals. b) Singular image for LV1 of reserve B-PLS showing negative voxel saliences (cool coloured regions). d) Singular image for LV1 of cognitive ability B-PLS showing negative voxel saliences. The scale represents the range of bootstrap ratio values thresholded at ± 3.28, p < 0.001. Activations are presented on template images of the lateral and medial surfaces of the left and right hemispheres of the brain using Multi-image Analysis GUI (Mango) software (http://ric.uthscsa.edu/mango/).

**Table 2:**
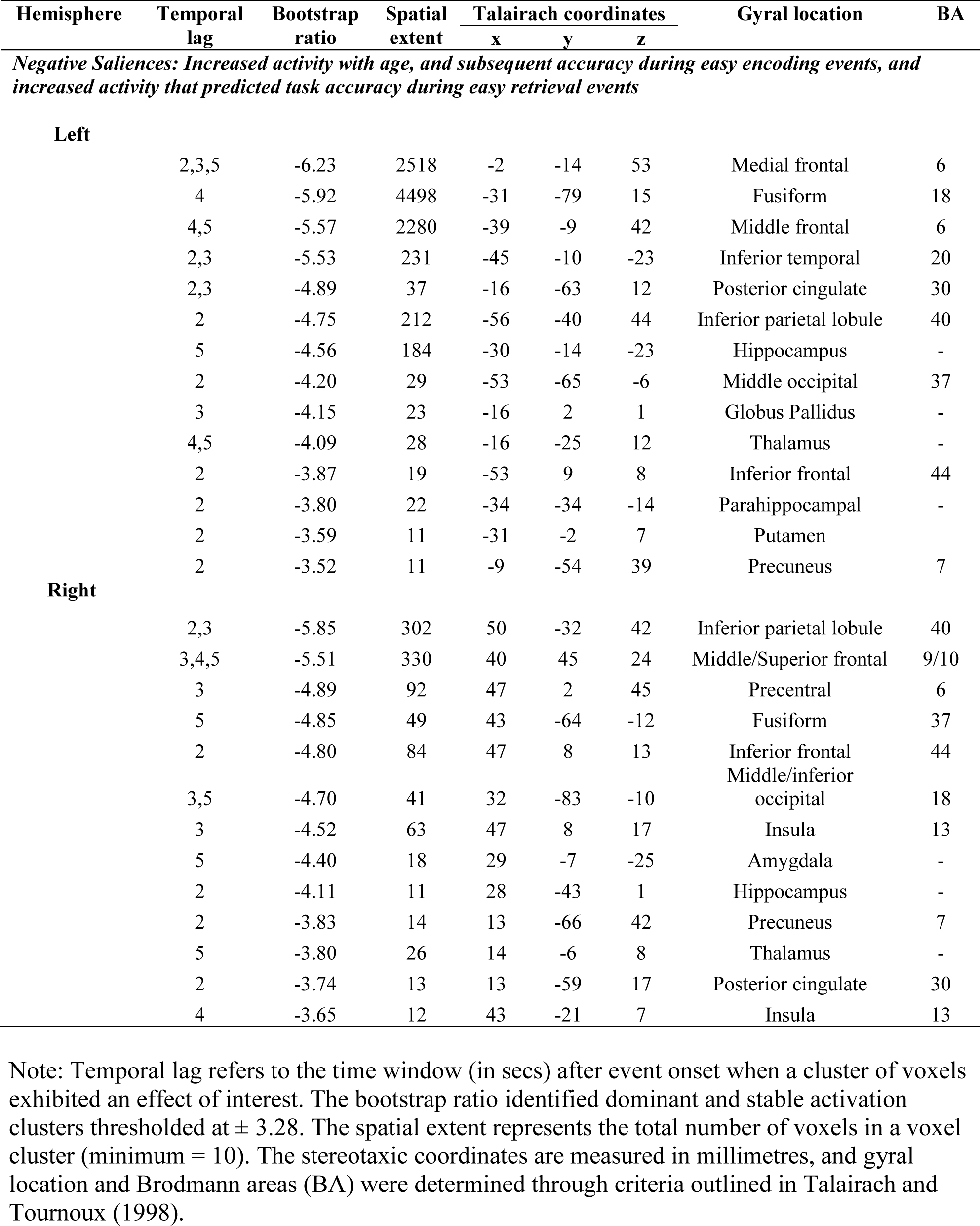
Local maxima for LV1 of reserve B-PLS: regions where activity correlated with age, and task accuracy.

Consistent with this interpretation, the post-hoc GLM for brain scores within easy encoding events against age, reserve, and task accuracy (R^2^ = 0.06, F (7, 300) = 2.83, p = .007) revealed significant main effects for age, accuracy (ps < 0.05), but not for reserve. On the other hand, post-hoc GLM for brain scores within easy retrieval events for all three variables (R^2^ = 0.05, F (7, 300) = 2.13, p = .04) revealed a significant main effect for task accuracy only (p < 0.001). There were no significant age*cognitive ability, age*accuracy or any other significant interactions revealed by the post-hoc tests. The negative salience brain regions represented in LV1 included: bilateral fusiform gyrus, medial frontal extending to ventrolateral PFC (BA 6/44), bilateral anterior PFC (BA 9/10), inferior parietal lobule (IPL), left frontal eye-fields (FEF: BA 8), left anterior temporal cortex, left hippocampus, and other regions (see Table 2).

LV2 accounted for 11.86% of the total cross-block covariance (p < 0.001) and primarily reflected a main effect of age identifying a whole-brain pattern of linear increases and decreases of activity with age across all encoding and retrieval events. The post-hoc GLM for brain scores against age, reserve, and task accuracy (R^2^ = 0.17, F (3,1228) = 82.73, p < .001) revealed a significant main effect for age across task conditions (p < 0.001), confirming this interpretation. The PLS correlation profile and the corresponding singular image are presented in Figures 2a and 2b respectively. Local maxima denoting positive and negative saliences for this LV are presented in Table 3. Age was positively correlated with activity in bilateral IPL, temporal cortex and right PHG (BA 28); and negatively correlated with activity in left fusiform cortex, posterior cingulate and thalamus. While some of the effects represented in this LV might be attributed to accuracy as shown in the PLS correlation profile, the effects observed resemble LV1 in our previous study (Ankudowich et al., 2016) and most strongly represent the effect of age on encoding and retrieval related activity across all tasks.

**Figure 2.**
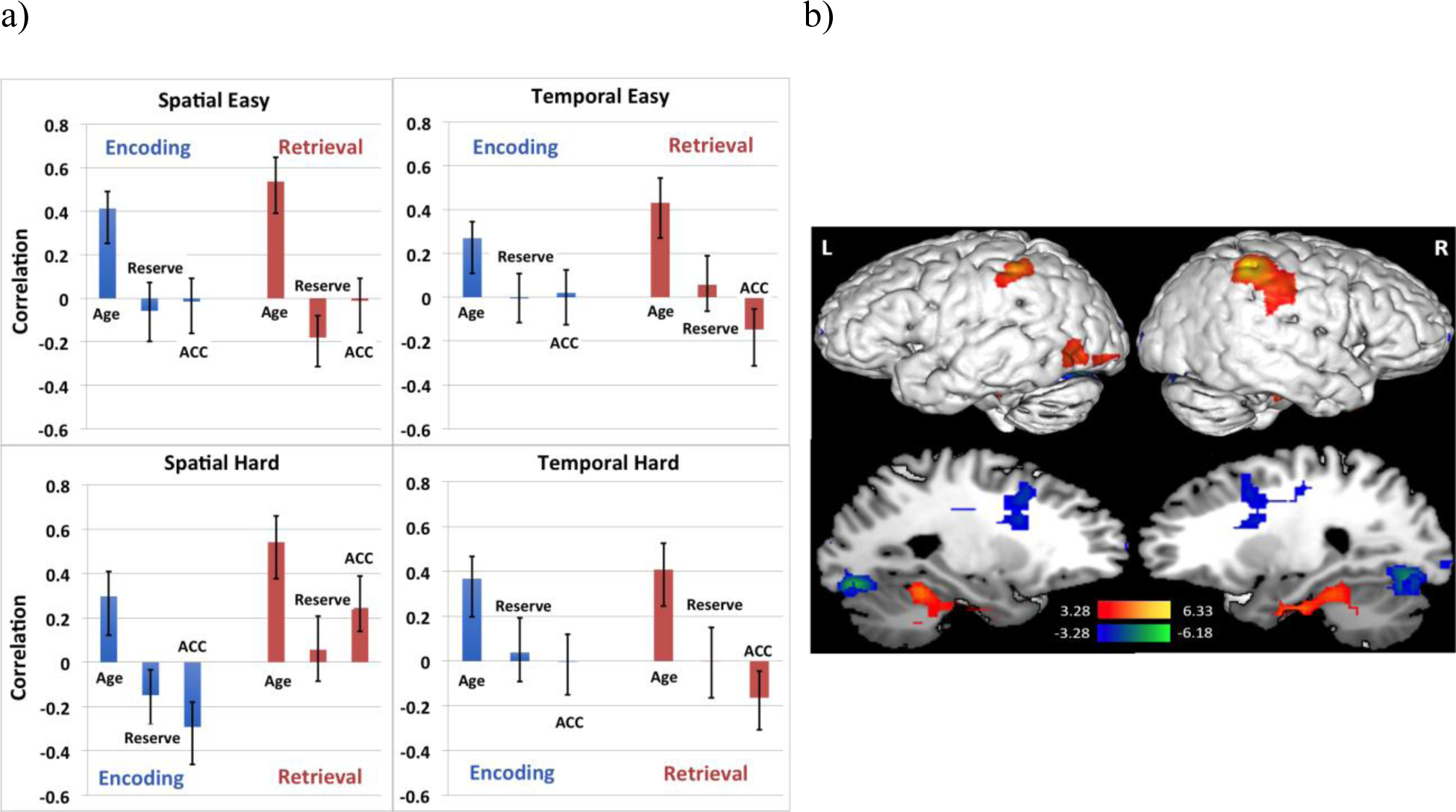
Brain-behaviour correlation profile and corresponding singular image for LV2 of reserve B-PLS analysis. a) LV2 brain-behaviour correlation profile for reserve B-PLS separated by task. The correlation profile indicated that activity in positive salience regions increased with age, and activity in negative salience regions decreased with age across tasks. ACC is short for accuracy. Error bars represent 95% confidence intervals. b) Singular image for LV2 of reserve B-PLS showing positive (warm coloured regions) and negative (cool coloured regions) voxel saliences. The scale represents the range of bootstrap ratio values thresholded at ± 3.28, p < 0.001. Activations are presented on template images of the lateral and medial surfaces of the left and right hemispheres of the brain using Multi-image Analysis GUI (Mango) software (http://ric.uthscsa.edu/mango/).

**Table 3:**
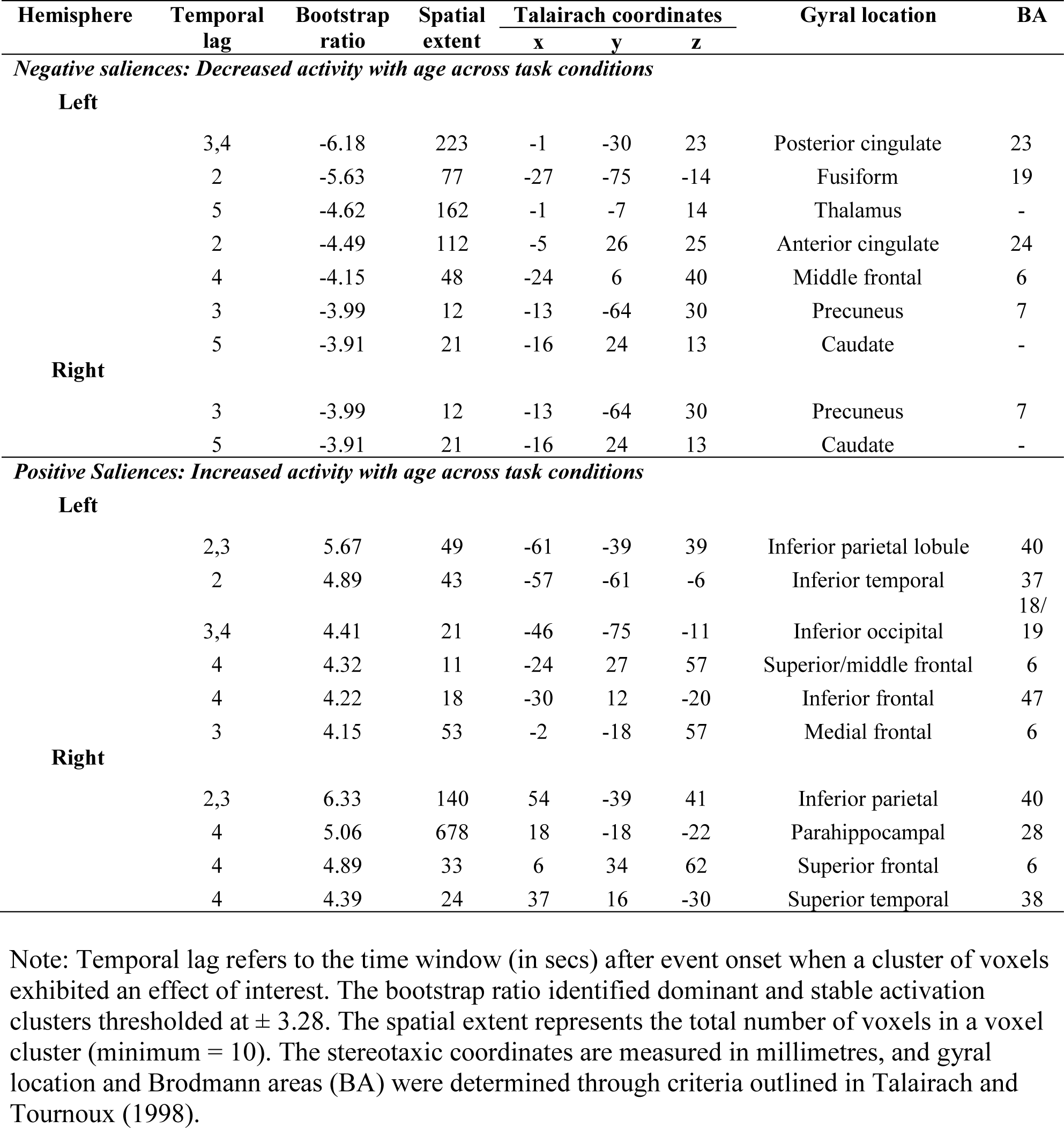
Local maxima for LV2 of reserve B-PLS: Regions where activity correlated with age across encoding and retrieval phases

LV3 accounted for 9.52% of the total cross-block covariance (p < 0.001). This LV identified brain regions that were differentially related to age during encoding and retrieval (age × phase effect). The local maxima of positive and negative voxel saliences are presented in Table 4. The PLS correlation profile and the corresponding singular image for LV3 are presented in Figures 3a and 3b respectively. Based on the PLS correlation profile, activity in positive salience brain regions (warm coloured regions in Figures 3b) were positively correlated with age during retrieval conditions (except TH retrieval), and negatively correlated with age at encoding. Positive salience brain regions included: bilateral hippocampus, IPL, putamen, superior temporal gyrys, and right ventrolateral PFC (BA 44). In contrast, activity in negative salience regions (blue coloured regions in Figures 3b) were positively correlated with age across all encoding conditions, and negatively correlated with age during retrieval. These regions included: bilateral fusiform gyrus, left postcentral gyrus, precuneus, and right precentral gyrus. Post-hoc GLMs for brain scores against age, reserve, and task accuracy at encoding (R^2^ = 0.06, F (3, 612) = 14.41, p < .001), and retrieval (R^2^ = 0.04, F (3, 612) = 9.77, p < .001), revealed significant main effects of age (ps < 0.01), but in opposite directions, mirroring the age × phase interaction outlined in the PLS correlation profile.

**Figure 3.**
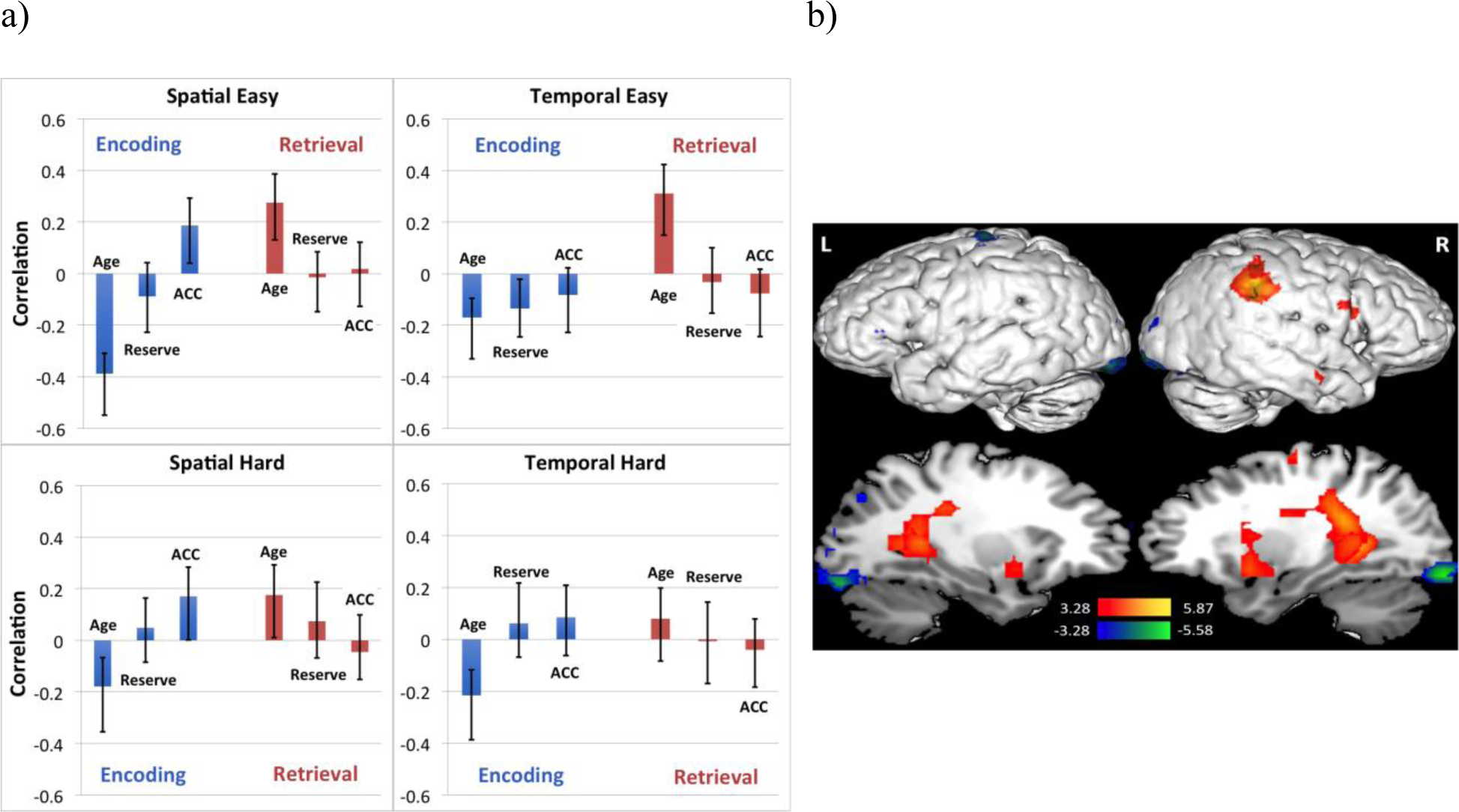
Brain-behaviour correlation profile and corresponding singular image for LV3 of reserve B-PLS analysis. a) LV3 brain-behaviour correlation profile for reserve B-PLS separated by task. The correlation profile indicated that activity in positive salience regions increased with age at retrieval and decreased with age at encoding. Activity in negative salience regions increased with age at encoding and decreased with age at retrieval. ACC is short for accuracy. Error bars represent 95% confidence intervals. b) Singular image for LV3 of reserve B-PLS showing positive voxel saliences (warm coloured regions) and negative voxel saliences (cool coloured regions). The scale represents the range of bootstrap ratio values thresholded at ± 3.28, p < 0.001. Activations are presented on template images of the lateral and medial surfaces of the left and right hemispheres of the brain using Multi-image Analysis GUI (Mango) software (http://ric.uthscsa.edu/mango/).

**Table 4:** Local maxima for LV3 of reserve B-PLS: regions where activity correlated with age differentially at encoding and retrieval.

| Hemisphere | Temporal lag | Bootstrap ratio | Spatial extent | Talairach coordinates |  |  | Gyral location | BA |
| --- | --- | --- | --- | --- | --- | --- | --- | --- |
|  |  |  |  | x | y | z |  |  |
| Negative Saliences: Increased activity with age at encoding and decreased at retrieval |  |  |  |  |  |  |  |  |
| Left | 2,3,4,5 | -5.39 | 107 | -39 | -34 | 62 | Postcentral gyrus | 1/2 |
|  | 2 | -4.80 | 20 | -20 | 55 | 18 | Middle frontal | 10 |
|  | 2,3 | -4.60 | 42 | -13 | -72 | 30 | Precuneus | 7/13 |
|  | 3 | -4.04 | 37 | -27 | -75 | -14 | Fusiform | 19 |
| Right | 3,4,5 | -5.58 | 90 | 25 | -87 | -14 | Fusiform | 18/37 |
|  | 4 | -5.11 | 39 | 46 | -11 | 58 | Precentral | 4/6 |
|  | Positive Saliences: Increased activity with age at retrieval and decreased at encoding |  |  |  |  |  |  |  |
| Left | 5 | 4.69 | 64 | -27 | -43 | 7 | Hippocampus | - |
|  | 2 | 4.40 | 42 | -23 | 7 | -6 | Putamen | - |
|  | 2 | 3.65 | 14 | -24 | -30 | 26 | Caudate | - |
| Right | 3,5 | 5.87 | 1017 | 50 | -35 | 31 | Inferior parietal lobule | 40 |
|  | 4 | 5.12 | 610 | 25 | -41 | 19 | Caudate Tail | - |
|  | 5 | 4.55 | 105 | 28 | -44 | 8 | Hippocampus | - |
|  | 2 | 4.42 | 84 | 21 | 10 | 2 | Putamen | - |
|  | 3 | 4.34 | 25 | 51 | 4 | 20 | Inferior fronal | 44 |
|  | 4 | 4.22 | 26 | 20 | -22 | 57 | Precentral | 4 |
|  | 2 | 4.16 | 11 | 55 | -8 | -10 | Superior temporal | 22 |
|  | 2 | 4.03 | 28 | 17 | -38 | 26 | Cingulate | 31 |
|  | 4 | 3.93 | 31 | 17 | -91 | 32 | Cuneus | 19 |
|  | 4 | 3.51 | 12 | 2 | -26 | 16 | Thalamus | - |

Note: Temporal lag refers to the time window (in secs) after event onset when a cluster of voxels exhibited an effect of interest. The bootstrap ratio identified dominant and stable activation clusters thresholded at $\pm 3.28$ . The spatial extent represents the total number of voxels in a voxel cluster (minimum = 10). The stereotaxic coordinates are measured in millimetres, and gyral location and Brodmann areas (BA) were determined through criteria outlined in Talairach and Tournoux (1998).

LV4 accounted for 6.94% of the total cross-block covariance (p < 0.001) and identified a pattern of brain activity that was mainly related to reserve across task conditions. The local maxima of negative and positive salience brain regions are presented in Table 5. Based on the PLS correlation profile (Figure 4a) and the corresponding singular image (Figure 4b), across all encoding and retrieval tasks (except SH encoding), activity in left superior temporal, caudate, and right cuneus increased with reserve. In contrast, activity in left dorsolateral PFC (BA 9) decreased with reserve. The post-hoc GLM model testing for brain scores against age, reserve, and task accuracy across task conditions (R^2^ = 0. 11, F (3,1228) = 52.87, p < .001) revealed a significant main effect for reserve (p < 0.001), confirming our interpretation.

**Figure 4.**
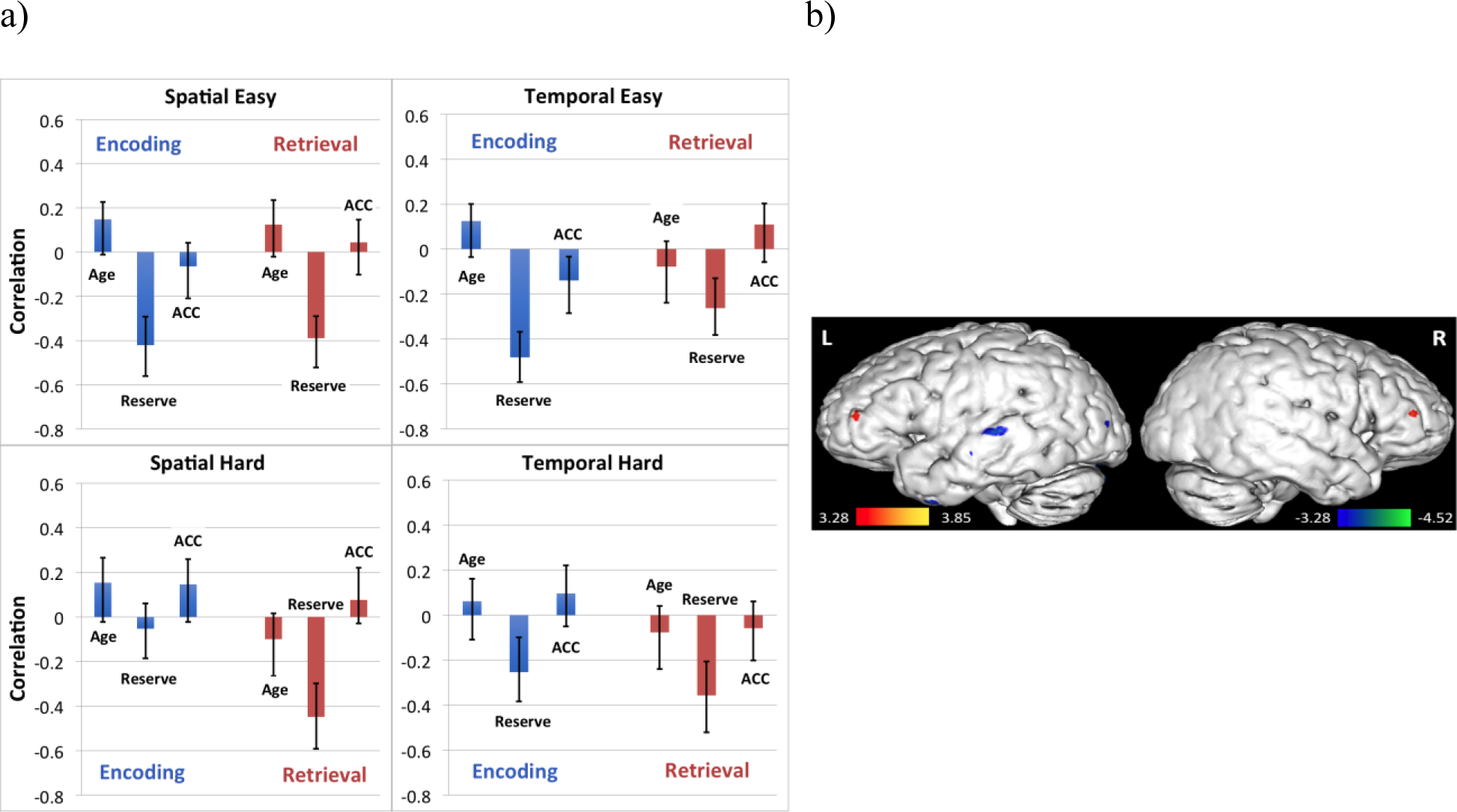
Brain-behaviour correlation profile and corresponding singular image for LV4 of reserve B-PLS. a) LV4 brain-behaviour correlation profile for reserve B-PLS separated by task. The correlation profile indicated that activity in negative salience regions increased with reserve across all task conditions (except for SH encoding), while activity in positive salience regions decreased with cognitive reserve across task conditions (except for SH encoding). ACC is short for accuracy. Error bars represent 95% confidence intervals. b) Singular image for LV4 showing negative voxel saliences (cool coloured regions). The scale represents the range of bootstrap ratio values thresholded at ± 3.28, p < 0.001. Peak activations were predominantly on the left and right lateral surfaces of the brain and were displayed using Multi-image Analysis GUI (Mango) software (http://ric.uthscsa.edu/mango/).

**Table 5:**
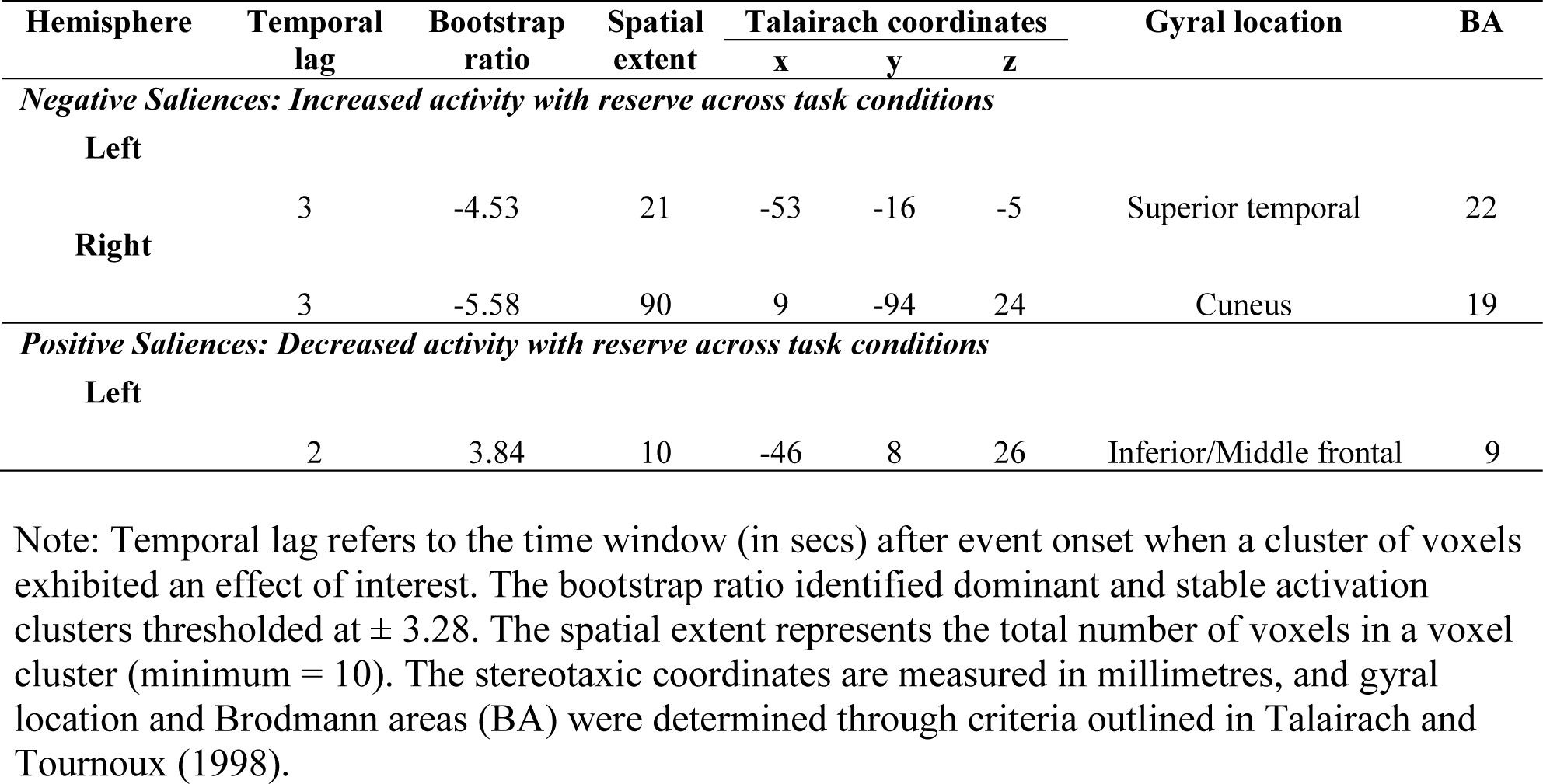
Local maxima for LV4 of reserve B-PLS: regions that were predominantly related to cognitive reserve across task conditions

The last significant LV (LV5) accounted for less than 5% of the total cross-block co-variance and showed minimal effects related to age, reserve, or accuracy, rending it uninterpretable. For this reason, LV5 will not be discussed further.

#### Post-hoc B-PLS: Age, Cognitive Ability and Accuracy

Given that our behavioural results indicated that our composite measure of cognitive ability predicted memory performance on our task fMRI paradigm (see above), whereas our proxy measure of reserve did not, we decided to conduct a second B-PLS analysis in which we used our composite measure of cognitive ability, in place of reserve, to explore if there were additional patterns of encoding- and retrieval-related brain activity that were positively associated with all three vectors of age, cognitive ability and memory performance. Such a result would indicate that cognitive ability may reflect aspects of reserve, which were not fully captured by our proxy measure of reserve, that in turn support memory performance in advanced age. In other words, this PLS analysis would allow us to determine if having higher cognitive ability supports the engagement of compensatory mechanisms with advancing age to support episodic memory processes. This approach is similar to that used in prior studies investigating the neural correlates of successful aging, and compensation, in high, compared to lower, preforming older adults (Cabeza et al., 2002; Duarte, Ranganath, Trujillo, & Knight, 2006; Glisky, Rubin, & Davidson, 2001; Nagel et al., 2009). Prior to running this B-PLS, we orthogonalized the cognitive ability measure to the age variable by obtaining its residual from a linear regression in which age was the predictor. This was done because age and cognitive ability were correlated (as stated above).

The B-PLS revealed four significant LVs that showed associations between the behavioural vectors of age, cognitive ability, and task accuracy; and whole brain patterns of event-related activity. The first LV accounted for 19.57 % of the cross-block covariance (p < 0.001) and revealed a pattern of activity that was similar to LV1 in the previous B-PLS analysis with reserve. The PLS correlation profile and the corresponding singular image are presented in Figures 1c and 1d respectively. Local maxima denoting negative saliences for this LV are presented in Table 6. Similar to LV1 in the first B-PLS analysis, activity in negative salience brain regions increased with age, predicted task accuracy during easy encoding events (SE and TE), and increased with task accuracy during easy retrieval events. However, in this analysis, activity in negative salience brain regions *also increased with cognitive ability during easy encoding events. C*onsistent with this interpretation, the post-hoc GLM model for brain scores within easy encoding events (SE, TE) against age, cognitive ability, and task accuracy (R^2^ = 0.09, F (7, 300) = 4.53, p < .001) revealed significant main effects for all 3 variables (ps < 0.05). On the other hand, post-hoc GLM for brain scores within easy retrieval events (SE, TE) for all three variables (R^2^ = 0.06, F (7, 300) = 2.881, p = .006) revealed a significant main effect only for task accuracy (p < 0.001). There were no significant age*cognitive ability, age*accuracy or any other significant interactions revealed by the post-hoc tests. The negative salience brain regions revealed in this LV are largely similar to those outlined in LV1 of the first analysis and included: bilateral fusiform gyrus (BA 18/37), medial frontal gyrus (BA 6), ventrolateral PFC (BA 44), right anterior PFC (BA 9/10), IPL (BA 40), left anterior temporal cortex (BA 20), bilateral hippocampus, and other regions (see Table 6). Therefore the first LV in both analyses identified a pattern of age-related increases in brain activity during easy memory tasks that supported retrieval performance. Although this pattern of activation was not related to our proxy measure of reserve (as indicated by our initial PLS), it was related to cognitive ability.

**Table 6:**
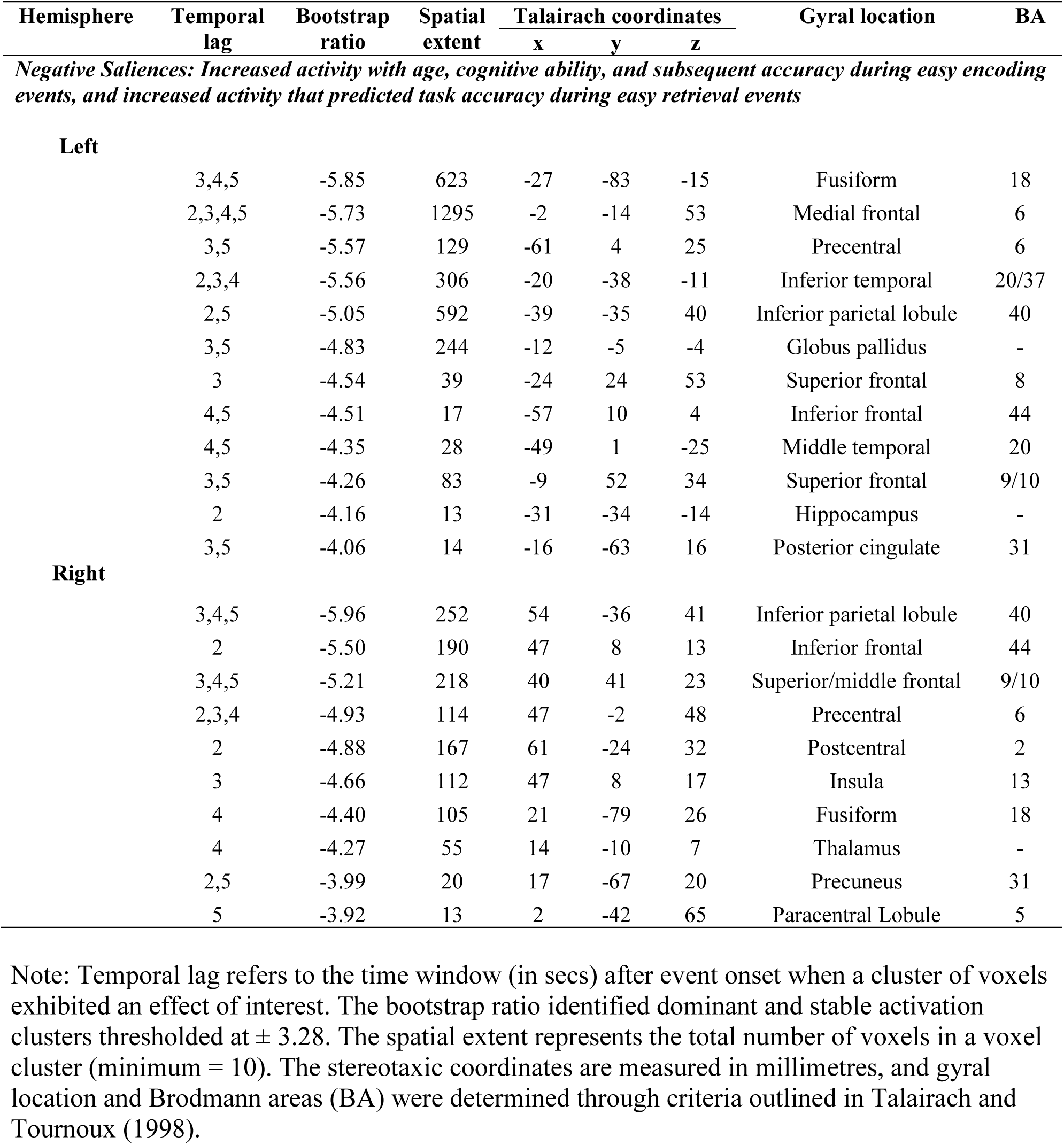
Local maxima for LV1 of cognitive ability B-PLS: regions where activity correlated with age, cognitive ability, and task accuracy.

| Hemisphere | Temporal lag | Bootstrap ratio | Spatial extent | Talairach coordinates |  |  | Gyral location | BA |
| --- | --- | --- | --- | --- | --- | --- | --- | --- |
|  |  |  |  | x | y | z |  |  |
| Negative Saliences: Increased activity with age, cognitive ability, and subsequent accuracy during easy encoding events, and increased activity that predicted task accuracy during easy retrieval events |  |  |  |  |  |  |  |  |
| Left | 3,4,5 | -5.85 | 623 | -27 | -83 | -15 | Fusiform | 18 |
|  | 2,3,4,5 | -5.73 | 1295 | -2 | -14 | 53 | Medial frontal | 6 |
|  | 3,5 | -5.57 | 129 | -61 | 4 | 25 | Precentral | 6 |
|  | 2,3,4 | -5.56 | 306 | -20 | -38 | -11 | Inferior temporal | 20/37 |
|  | 2,5 | -5.05 | 592 | -39 | -35 | 40 | Inferior parietal lobule | 40 |
|  | 3,5 | -4.83 | 244 | -12 | -5 | -4 | Globus pallidus | - |
|  | 3 | -4.54 | 39 | -24 | 24 | 53 | Superior frontal | 8 |
|  | 4,5 | -4.51 | 17 | -57 | 10 | 4 | Inferior frontal | 44 |
|  | 4,5 | -4.35 | 28 | -49 | 1 | -25 | Middle temporal | 20 |
|  | 3,5 | -4.26 | 83 | -9 | 52 | 34 | Superior frontal | 9/10 |
|  | 2 | -4.16 | 13 | -31 | -34 | -14 | Hippocampus | - |
|  | 3,5 | -4.06 | 14 | -16 | -63 | 16 | Posterior cingulate | 31 |
| Right | 3,4,5 | -5.96 | 252 | 54 | -36 | 41 | Inferior parietal lobule | 40 |
|  | 2 | -5.50 | 190 | 47 | 8 | 13 | Inferior frontal | 44 |
|  | 3,4,5 | -5.21 | 218 | 40 | 41 | 23 | Superior/middle frontal | 9/10 |
|  | 2,3,4 | -4.93 | 114 | 47 | -2 | 48 | Precentral | 6 |
|  | 2 | -4.88 | 167 | 61 | -24 | 32 | Postcentral | 2 |
|  | 3 | -4.66 | 112 | 47 | 8 | 17 | Insula | 13 |
|  | 4 | -4.40 | 105 | 21 | -79 | 26 | Fusiform | 18 |
|  | 4 | -4.27 | 55 | 14 | -10 | 7 | Thalamus | - |
|  | 2,5 | -3.99 | 20 | 17 | -67 | 20 | Precuneus | 31 |
|  | 5 | -3.92 | 13 | 2 | -42 | 65 | Paracentral Lobule | 5 |

Note: Temporal lag refers to the time window (in secs) after event onset when a cluster of voxels exhibited an effect of interest. The bootstrap ratio identified dominant and stable activation clusters thresholded at $\pm 3.28$ . The spatial extent represents the total number of voxels in a voxel cluster (minimum = 10). The stereotaxic coordinates are measured in millimetres, and gyrus location and Brodmann areas (BA) were determined through criteria outlined in Talairach and Tournoux (1998).

Subsequent LVs 2, 3 replicated the LV2 (main effect of age; accounted for 12.27% of the total cross-block covariance) and LV3 (age*phase interaction; accounted for 10.15% of the total cross-block covariance) results obtained in our first PLS analyses and therefore are not discussed further here. These results are presented in supplementary Tables 1 and 2 and supplementary Figures 1 and 2, respectively. The fourth LV accounted for 6.41% of the total cross-block covariance (p < 0.001) and identified a pattern of brain activity that was mainly related to cognitive ability, independent of age. Only negative salience brain regions from this LV survived our spatial threshold cut-off of 10 contiguous voxels (p < 0.001), and the local maxima of those regions are presented in Table 7. Based on the PLS correlation profile (Figure 5a) and the corresponding singular image (Figure 5b), activity in left middle frontal gyrus, left fusiform gyrus, right precentral gyrus, and FEF (BA 8), was negativity correlated with cognitive ability during all retrieval conditions (and encoding SE). Consistent with this interpretation, the post-hoc GLM model testing for brain scores at retrieval against age, cognitive ability, and task accuracy (R^2^ = 0.09, F (3, 612) = 22.56, p < .001) revealed a significant main effect for cognitive ability (p < 0.001).

**Figure 5.**
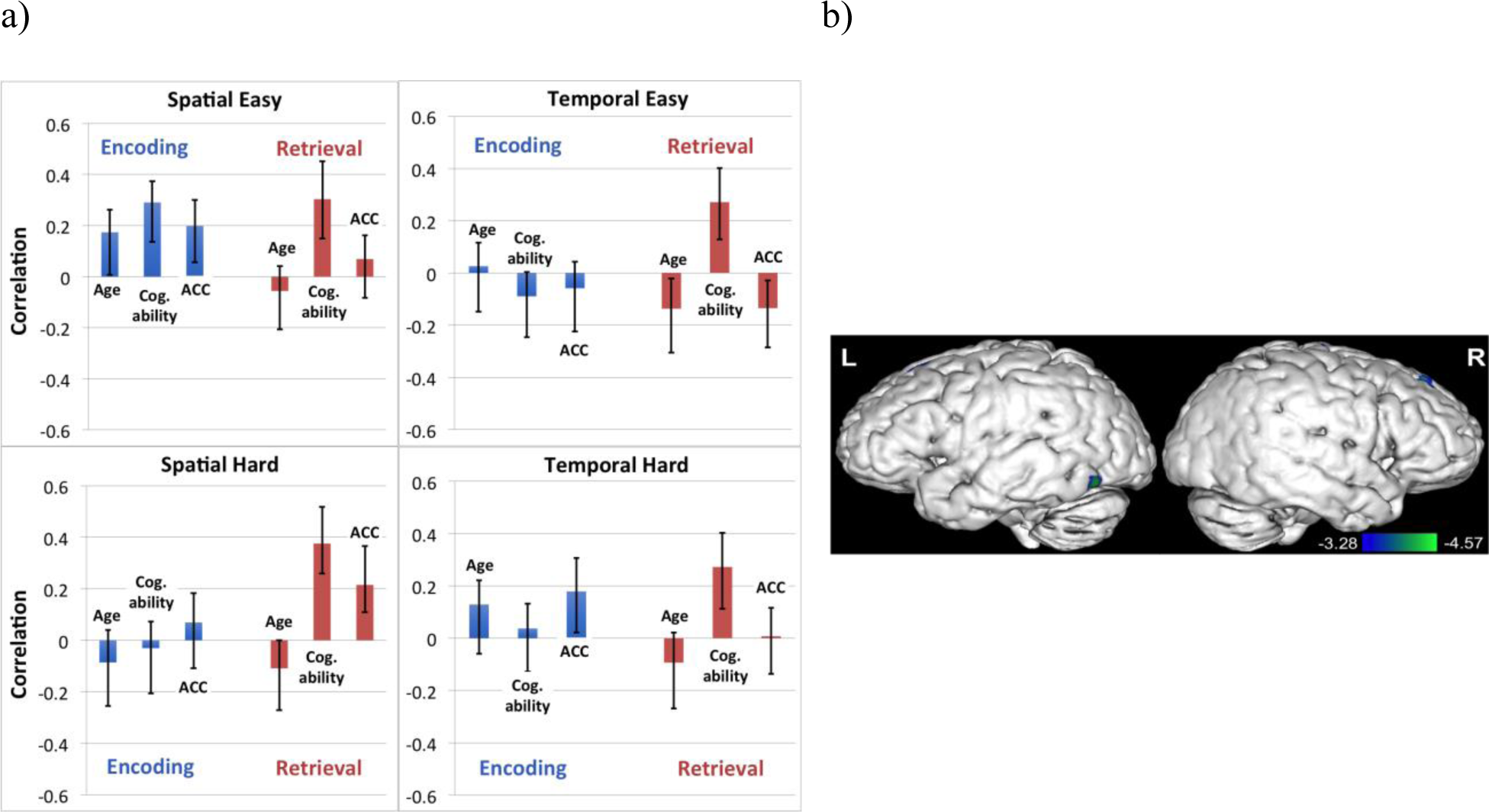
Brain-behaviour correlation profile and corresponding singular image for LV4 of cognitive ability B-PLS. a) LV4 brain-behaviour correlation profile for cognitive ability B-PLS separated by task. The correlation profile indicated that activity in negative salience regions decreased with cognitive ability across all retrieval events. ACC is short for accuracy, and Cog. ability is short for cognitive ability. Error bars represent 95% confidence intervals. b) Singular image for LV4 showing negative voxel saliences (cool coloured regions). The scale represents the range of bootstrap ratio values thresholded at ± 3.28, p < 0.001. Peak activations were predominantly on the left and right lateral surfaces of the brain and were displayed using Multi-image Analysis GUI (Mango) software (http://ric.uthscsa.edu/mango/).

**Table 7:**
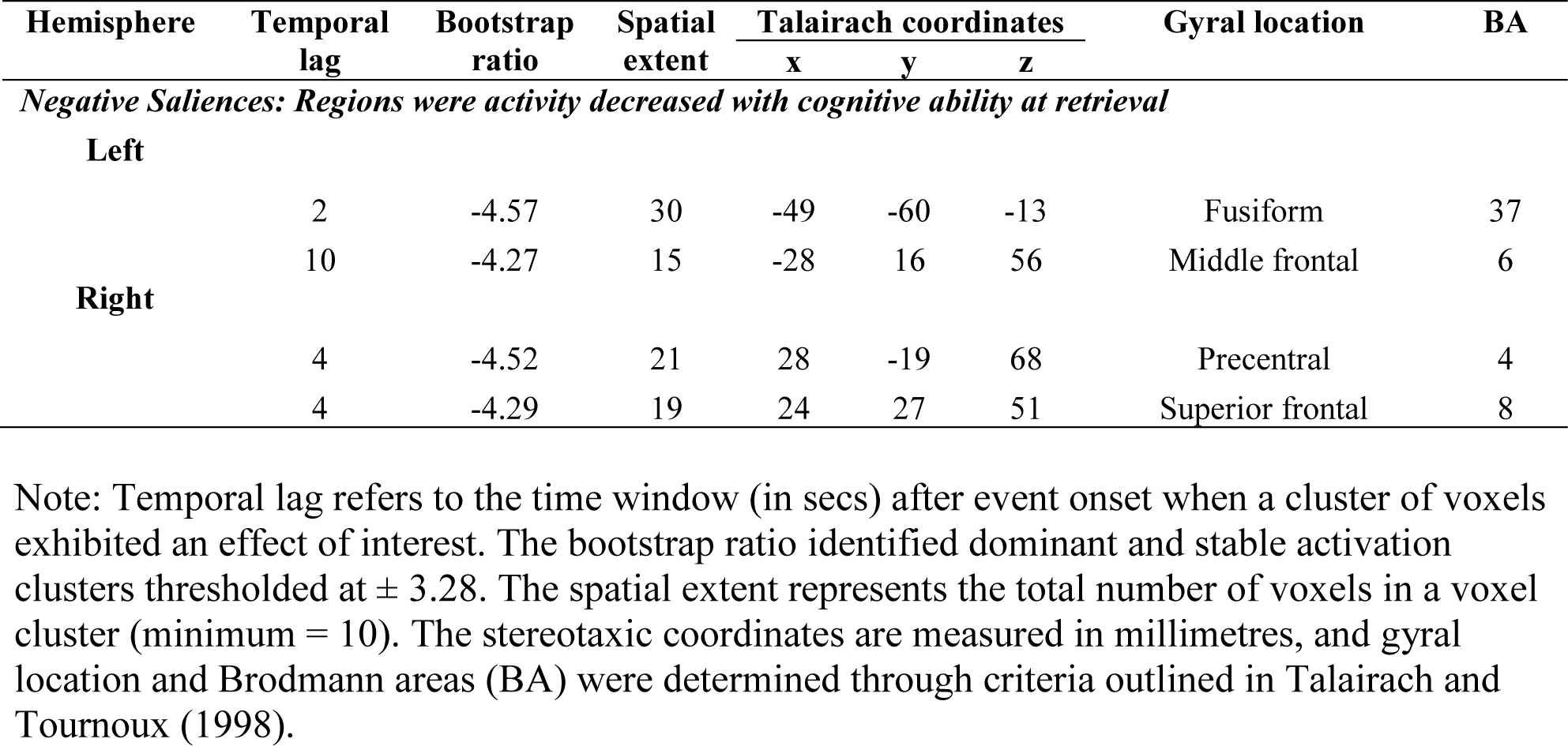
Local maxima for LV4 of cognitive ability B-PLS: regions that were predominantly related to cognitive ability at retrieval

| Hemisphere | Temporal lag | Bootstrap ratio | Spatial extent | Talairach coordinates |  |  | Gyral location | BA |
| --- | --- | --- | --- | --- | --- | --- | --- | --- |
|  |  |  |  | x | y | z |  |  |
| Negative Saliences: Regions were activity decreased with cognitive ability at retrieval |  |  |  |  |  |  |  |  |
| Left |  |  |  |  |  |  |  |  |
|  | 2 | -4.57 | 30 | -49 | -60 | -13 | Fusiform | 37 |
|  | 10 | -4.27 | 15 | -28 | 16 | 56 | Middle frontal | 6 |
| Right |  |  |  |  |  |  |  |  |
|  | 4 | -4.52 | 21 | 28 | -19 | 68 | Precentral | 4 |
|  | 4 | -4.29 | 19 | 24 | 27 | 51 | Superior frontal | 8 |

Note: Temporal lag refers to the time window (in secs) after event onset when a cluster of voxels exhibited an effect of interest. The bootstrap ratio identified dominant and stable activation clusters thresholded at $\pm 3.28$ . The spatial extent represents the total number of voxels in a voxel cluster (minimum = 10). The stereotaxic coordinates are measured in millimetres, and gyrus location and Brodmann areas (BA) were determined through criteria outlined in Talairach and Tournoux (1998).

## 4. Discussion

In the current adult lifespan task fMRI study, we tested the behavioural hypothesis that our proxy measure of reserve moderated the effect of age on our composite measure of cognitive ability and measures of episodic memory function obtained from task fMRI. We also tested the neural hypothesis that greater reserve would be related to decreased activity (greater neural efficiency) in lateral PFC and parietal regions (Stern et al., 2018), and increased activity (greater neural capacity / flexibility) in additional PFC regions, medial temporal and ventral visual regions, across the adult lifespan; and this in turn would related to better episodic memory ability (Ankudowich et al., 2017; Cabeza et al., 2002; Springer et al., 2005). To test these hypotheses, we created a proxy measure of reserve that included levels of educational attainment and premorbid intelligence (NART-I.Q.), and a composite measure for cognitive ability that included items from a neuropsychological test battery reflecting verbal memory ability (CVLT) and executive function (D-KEFS, and WCST). We then conducted a B-PLS analysis to examine how age, reserve, and retrieval accuracy correlated with brain activity during easy and hard, spatial and temporal context memory encoding and retrieval tasks. Based on our behavioural findings (discuss below), we also conducted a second, exploratory, B-PLS analysis to examine how age, cognitive ability, and task accuracy were associated with brain activity patterns across task conditions.

The behavioural regression analysis revealed that greater levels of reserve, as measured by education and I.Q., did not predict cognitive ability or task-fMRI context memory performance in our adult lifespan sample. Nor did higher reserve moderate the negative effect of age on cognitive ability and context memory retrieval accuracy. However, our regression and ANOVA results showed the typical pattern of age-related decrements in cognitive ability and context memory retrieval accuracy across all tasks. As expected, YA outperformed middle-aged and OA, and middle-aged adults performed better than OA. This is consistent with prior findings suggesting that context memory declines begin as early as mid-life (Cansino, 2009), and persist into older adulthood (Simons, Dodson, Bell, & Schacter, 2004; Wegesin et al., 2000). Participants in the current study also showed higher accuracy on easy relative to hard tasks, implying that increasing encoding load increased task difficulty.

The fact that we did not find a significant association between reserve and cognitive ability, memory performance and age-related memory decline is consistent with previous findings (Zahodne et al., 2011). However, other cross-sectional studies have shown a positive association between reserve proxies and episodic memory performance across age (Angel et al., 2010; Corral, Rodriguez, Amenedo, Sanchez, & Diaz, 2006; Lachman et al., 2010). The inconsistency between the current results and the aforementioned studies may partly be due to methodological differences. For example, reserve has been examined through the scope of premorbid I.Q. (Corral et al., 2006), or years of education (Angel et al., 2010; Lachman et al., 2010) separately, while we created a composite measure of reserve combining both measures. Furthermore, Corral et al. (2006) and Angel et al. (2010) categorized participants into high vs. low reserve groups, unlike the current study which examined reserve and episodic memory performance as continuous variables. It is also likely that our strict inclusion criteria may have contributed to skewing our sample towards individuals with higher levels of education. Our current sample included participants with years of education ranging from 11-20 years, which suggests the sample consisted of relatively highly educated individuals. In contrast the study by Angel et al. (2010) included a sample with years of education ranging from 8-17 years. It is possible that inclusion of more individuals with lower levels of education may help adequately capture the positive relationship between reserve and episodic memory performance. Nonetheless, our post-hoc regression analyses indicated that higher levels of cognitive ability was associated with better context memory performance, suggesting a potential protective role of other aspects of reserve on memory in advanced age not fully captured by our proxy measure of NART-I.Q. and years of education composite.

In relation to the fMRI findings, both B-PLS analyses identified similar effects linking brain activity and age across all task conditions (LV 2 in both analyses), and brain activity and age-by-phase (encoding and retrieval) interactions (LV 3 in both analyses). We have observed similar results in our prior analyses of a subset of this dataset (Ankudowich 2016, 2017) and have interpreted these results in our prior publications. In general, findings from LVs 2 and 3 are largely consistent with observations from previous fMRI studies of episodic memory across the adult lifespan and show that aging may be related to increases in lateral occipital-temporal, medial temporal and parietal regions activity, and decreases in fusiform activity (e.g., Grady, Springer, Hongwanishkul, McIntosh, & Winocur, 2006; Kennedy et al., 2012).

Additionally, the first LV in both B-PLS analyses revealed that activity in bilateral ventrolateral and right dorsolateral PFC, bilateral MTL (including the hippocampus), inferior parietal, precuneus and ventral occipito-temporal activity was positively correlated with age and subsequent memory. Interestingly, these age-related increases in activity were also associated with higher cognitive ability, but not with higher reserve as a function of task demands. This pattern demonstrated that task-specific compensatory responses related to enhanced neural capacity in the aging brain are not strongly associated with higher scores on reserve proxies of education and I.Q., but may be modulated by individual differences in cognitive ability. This finding dovetails with our behavioural results showing a positive association between task accuracy and cognitive ability, but not reserve. Therefore, based on our behavioural and fMRI findings, reserve (as indexed by education and premorbid I.Q.) does not moderate the effect of age on cognitive ability and measures of episodic memory function, nor does it relate to task specific compensatory responses with age in the current study. In addition, we observed distinct activity in the left medial-superior PFC in the first LV of the cognitive ability-PLS analysis.

Finally, there was a significant LV in each of the B-PLS analyses that identified brain regions in which task-related activity was correlated with reserve (LV4 of the reserve-PLS) and with cognitive ability (LV4 in the cognitive ability-PLS), but not age and retrieval accuracy. This suggest that individual differences in reserve and cognitive ability was related to unique patterns of brain activity during context memory encoding and retrieval, which did not directly support memory performance and was not impacted by aging. Below, we discuss these results, and results from the first LV in greater detail.

### Patterns of activity modulated by reserve

Across both B-PLS analyses, the LV explaining the largest amount of variance revealed a pattern of event-related activity that increased with age and predicted task accuracy during easy encoding trials. However, this pattern was associated with individual differences in cognitive ability (discussed below) and not reserve. This result contradicts our prediction that increased reserve would be directly related to different patterns of brain activity during easy, compared to difficulty memory tasks, which in turn would support memory performance. Indeed, the only LV that demonstrated a reliable whole-brain activity pattern related to our composite measure of reserve (i.e., education and I.Q.) was LV4. This LV highlighted increases in left superior temporal and cuneus activity, paired with decreases in left inferior frontal activity that were correlated with higher reserve across task conditions. It has been suggested that the neural implementation of reserve manifests as a domain-general pattern that is expressed across a variety of cognitive tasks and that the degree of expression of this pattern would correlate with reserve proxies like education and I.Q. (Cabeza et al., 2018). The current findings are consistent with this notion and demonstrate that reserve (as indexed by education and premorbid I.Q.) was associated with linear increases and decreases in brain activity across easy and hard spatial/temporal context memory, both during encoding and retrieval. Stern and colleagues (2018) used a multivariate analysis approach to identify a task-general pattern of activity that correlates with a proxy of reserve (NART-I.Q.) in individuals aged 20-80 years old. Across 12 different cognitive tasks including episodic memory, they found that activity in several regions including cerebellum, medial frontal and superior temporal regions increased with reserve, while activity in inferior frontal and parietal regions decreased with reserve, consistent with the current results. The fact that the task-general pattern observed in LV4 correlated with reserve and not with age suggests that this pattern is expressed as a function of reserve, regardless of age-related cognitive change. Similar to arguments made by Stern et al., (2018), we propose that this pattern of activity related to reserve is available throughout the adult lifespan and may set up individuals to deal with age-related changes as they occur in old age. This suggestion is also in line with the concept of ‘neural reserve’ which posits that individual differences in brain networks modulated by reserve may allow some individuals to cope with the disruption related to age or brain pathology (Stern, 2009). Further investigation is required to test this hypothesis.

### Patterns of activity modulated by cognitive ability

The first LV in both B-PLS analyses identified a set of distributed negative salience brain regions including bilateral ventral occipital-temporal cortices, IPL, precuneus, PFC, and MTL. Activity in these regions increased with age during easy context encoding tasks and was positively correlated with subsequent memory on these tasks. Interestingly, these age-related increases were also modulated by cognitive ability (but not by reserve). Our findings suggest that compensatory responses in older age are not modulated by the reserve proxies of education and I.Q., but are rather associated with individual differences in cognitive ability. During retrieval, activity in those regions also appeared to support performance on easier versions of the task, albeit not in association with age or cognitive ability. This implies that at retrieval, re-activation of the same network of regions initially recruited during encoding supports memory accuracy for the same event types, adding to the rich body of literature arguing that successful recollection hinges on reinstatement or recapitulation of the cognitive and/or neural processes engaged during memory encoding (Buckner & Wheeler, 2001; Rugg, Johnson, Park, & Uncapher, 2008; Tulving, Voi, Routh, & Loftus, 1983; Waldhauser, Braun, & Hanslmayr, 2016; Wheeler, Petersen, & Buckner, 2000). Interestingly, we only observed this pattern of association during easy memory tasks. We did not identify an LV that related cognitive ability, age and performance during difficult memory tasks. This suggests that having higher cognitive ability is positively correlated with functional compensatory activity in lateral and medial prefrontal, parietal, medial temporal and occipito-temporal regions, but only during less demanding easy memory tasks.

As stated above, the effects in LV1 were reliably observed in ventral occipital cortex and lateral temporal cortices. We have previously shown that age was positively associated with ventral occipital and fusiform cortex at encoding, but this was not correlated to subsequent memory (Ankudowich et al., 2017). Instead, ventral occipital and fusiform activity at retrieval was positively correlated with memory performance; however, older adults exhibited reduced activity in these ventral occipital cortex regions at retrieval. We also found that age was positively correlated with lateral temporal activity across context encoding and retrieval tasks, but this was not positively correlated with memory performance. Interestingly, LV1 in the current study revealed a novel finding that cognitive ability was related to an age-related increase in both ventral occipital and lateral temporal activation during easy encoding tasks, and this was related to better subsequent memory. These ventral occipital cortical regions, including fusiform cortex, have consistently been shown to be related to face processing and recognition (Haxby, Hoffman, & Gobbini, 2000; Loffler, Yourganov, Wilkinson, & Wilson, 2005; Rotshtein, Henson, Treves, Driver, & Dolan, 2005). In contrast, previous neuroimaging studies of memory have shown that activity in lateral temporal regions support more generalized semantic/conceptual processing of visual stimuli (Binder & Desai, 2011; Menon, Boyett-Anderson, Schatzberg, & Reiss, 2002). Therefore, our current results suggest that individuals with higher cognitive ability may be better positioned to engage a variety of visual encoding strategies with advanced age, and that this benefits successful memory encoding and retrieval.

Individuals with higher cognitive ability also activated left anterior hippocampus more with advanced age at encoding. The anterior hippocampus has been shown to be more active during episodic encoding, compared to retrieval (Kim, 2015; Lepage, Habib, & Tulving, 1998) and to contribute to relational processes (Davachi, 2006) and conceptual encoding requiring the integration of a variety of perceptual, emotional and semantic information (Zeidman & Maguire, 2016). We have previously shown that larger anterior hippocampal volumes were associated with better spatial and temporal context memory in young adults (Rajah et al., 2010). We also found that age-related reduction in anterior hippocampal volumes was associated with increased encoding activity in occipital, lateral temporal and PFC, which was also associated with better subsequent memory for easier, compared to harder, context memory task in older adults (Maillet & Rajah, 2013). The current results corroborate our prior findings and indicate that older adults with higher cognitive ability were better able to co-activate anterior hippocampal, occipital-temporal and PFC regions (discussed below) at encoding, which benefitted subsequent memory during easy tasks.

LV1 also identified activations in precuneus and inferior parietal regions (BA 7 and BA 40 respectively), ventrolateral PFC (BA 44) and dorsal PFC (BA8; frontal eye fields). Evidence from the attention literature points to the presence of two functionally distinct attention systems in the human brain. A dorsal fronto-parietal system involving superior parietal (precuneus) regions and frontal eye-fields, which is thought to be involved in top-down allocation of attentional resources to locations or different features; and a ventral fronto-parietal system involving inferior parietal regions and ventrolateral PFC, which is thought to be involved in stimulus-driven, bottom-up shifts in attentional focus. Whether age- and cognitive ability-related increases in fronto-parietal activity observed in LV1 reflected supervisory top-down, or stimulus-driven bottom up attentional processes cannot be discerned from the current results. Nevertheless, recent evidence suggests that these two systems do not operate independently and interact to allow for the flexible control of attention in response to current task demands (Vossel, Geng, & Fink, 2014). We also observed distinct activity in the left medial-superior PFC (BA 9/10) that was unique to the cognitive ability B-PLS in that activity in this region increased with age, cognitive ability, and predicted successful task accuracy during easy encoding events. We have previously shown that activity in the left superior PFC during encoding and retrieval correlates with better spatial and context memory performance, but decreases with age (Ankudowich et al., 2017). However, our results reveal that despite the overall age-related decrease in activity in this region, older individuals with higher cognitive ability are able to maintain high levels of function in superior PFC supporting context memory performance. Taken together, our findings indicate that older adults with greater cognitive ability may be better able to recruit frontoparietal cognitive control processes to modulate the engagement of the aforementioned visual and mnemonic strategies to benefit memory performance during easy context memory tasks.

While the pattern of results in LV1 identified brain regions that supported task performance with advanced age, age-related deficits were still apparent. In the current study, since the pattern of age-related over-recruitment observed in LV1 was associated with better task performance, it can be regarded as a form of functional compensation (Cabeza & Dennis, 2013; Grady, 2008; Rajah & D’Esposito, 2005) to cope with task demands even at easier versions of the context memory task. At higher levels of task difficulty (i.e., SH and TH), the set of brain regions that aided performance on easy tasks were no longer associated with age, cognitive ability, or performance. It is conceivable that easier versions of the task in the current study placed higher cognitive demands as a function of age, and with increased age and cognitive ability, individuals were better able to engage occipito-temporal, hippocampal and frontoparietal brain regions to support task performance. This interpretation is consistent with the CRUNCH model (Reuter-Lorenz & Cappell, 2008; Reuter-Lorenz & Lustig, 2005), which suggests that load-sensitive, task-related brain regions are recruited in older age at lower levels of task demands compared to YA who may recruit those regions at higher levels of task demand. The fact that age-related decrements in accuracy were observed in both spatial easy (F (2,151) = 6.88, p < 0.001, η2 = 0.083), and temporal easy (F (2,151) = 11.87, p < 0.001, η2 = 0.136) tasks lends further support to this interpretation. This finding is also consistent with the notion of neural capacity, which posits that individuals with higher reserve are able to maximize recruitment of task-related regions under increasing demands to support task performance (Barulli & Stern, 2013). Although this pattern was associated with cognitive ability and not with our proxy measure of reserve (education and I.Q.), cognitive ability may in fact be considered an outcome of reserve and may reflect other reserve factors not accounted for by education or I.Q. Therefore, findings from LV1 suggest that demand-related compensatory responses observed with increased age during memory encoding are modulated by individual differences in cognitive ability.

The last significant (LV4) from the cognitive ability B-PLS identified a small subset of brain regions, including left fusiform and motor cortices, that showed attenuations in activity at retrieval associated with higher cognitive ability. This observation is akin to the hypothesis that higher reserve relates to higher neural efficiency as indexed by lower activity in task-related brain regions (Barulli & Stern, 2013; Steffener & Stern, 2012; Stern, 2009). However, since the attenuations in activity were observed at both lower and higher levels of task difficulty, it is unclear to us whether findings from this LV does in fact reflect enhanced neural efficiency as a function of cognitive ability (or reserve factors apart from education and I.Q.), and whether this finding would generalize across studies.

Besides the overall patterns of activity associated with cognitive ability discussed above, it is important to point out that distinct regions of the medial temporal lobes were identified across LVs 1 through 3, and activity in those regions were deferentially related to cognitive ability. Increased age-related activity in left anterior hippocampus was observed in LV1, and was positively correlated with cognitive ability and subsequent memory. Age-related increases in right PHG was observed at encoding and retrieval, and age-related increases in bilateral posterior HC were observed at retrieval (LV3). Interestingly, age-related increases in right PHG activity at encoding was negatively related to cognitive ability, but age-related increases in bilateral posterior hippocampus and anterior hippocampus activity were positively related to cognitive ability. Together, these observations suggest that individuals with higher vs. lower cognitive ability may exhibit age-related differences in hyperactivation within the MTL. Age-related increases in hippocampal activity may be positively related to cognitive ability, and this suggests that older adults may exhibit different patterns of MTL hyper activation as a function of cognitive ability during encoding vs. retrieval. There has been significant interest in the cognitive neuroscience of aging and dementia fields on whether age-related increases in MTL activation reflect compensation in healthy aging (Cabeza et al., 2018), or whether these ‘hyperactivations’ may reflect a neurotoxic response in the aging brain which may be an early sign of pathological aging (Bookheimer et al., 2000; Fleisher et al., 2009; Quiroz et al., 2010). Our results suggest that individuals with higher cognitive ability exhibit increased hippocampal activity with age, and that this increase at encoding may reflect compensation. In contrast, it is less clear whether the increased parahippocampal activation in lower reserve adults in advanced age is detrimental to performance or not. This is an important topic for future research, given the importance of hippocampal function in memory and the rich literature implicating age-related difference in MTL function in both healthy and pathological aging.

## Conclusions

In sum, the current study provided novel evidence that task-specific, age-related compensatory responses in inferior parietal, ventral visual, and PFC during context memory encoding are modulated by individual differences in cognitive ability, but not by a composite proxy measure of reserve (as indexed by education and I.Q.). In contrast, higher reserve is associated with task-general responses in superior temporal, occipital, and inferior frontal regions. Although higher reserve was not directly associated with increased compensation in the current study, the relationship between compensation and reserve appears to be complex and task-dependent. Higher cognitive ability in old age is assumed to be an outcome of accumulated reserve (Stern et al., 2018), and was associated with age-related functional compensation in the current study as a function of task demands. Therefore, it is reasonable to expect that lifestyle factors indexing cognitive reserve such as education and I.Q. would also be associated with functional compensation as a function of age. This was not supported by our current findings. It is possible that the proxy measures of education and I.Q. utilized in the current study did not fully capture accumulated reserve, and that other sociobehavioural factors such as occupational complexity, social interaction, leisure, physical activity and other protective factors should be considered when examining the impact of cognitive reserve on brain activity. An important topic for future investigation would be examining the various factors that contribute to the development of reserve, and how these factors individually, or collectively relate to cognitive ability and alterations in brain activity to cope with structural and functional changes in the aging brain.

---

**Supplementary Figure 1.**
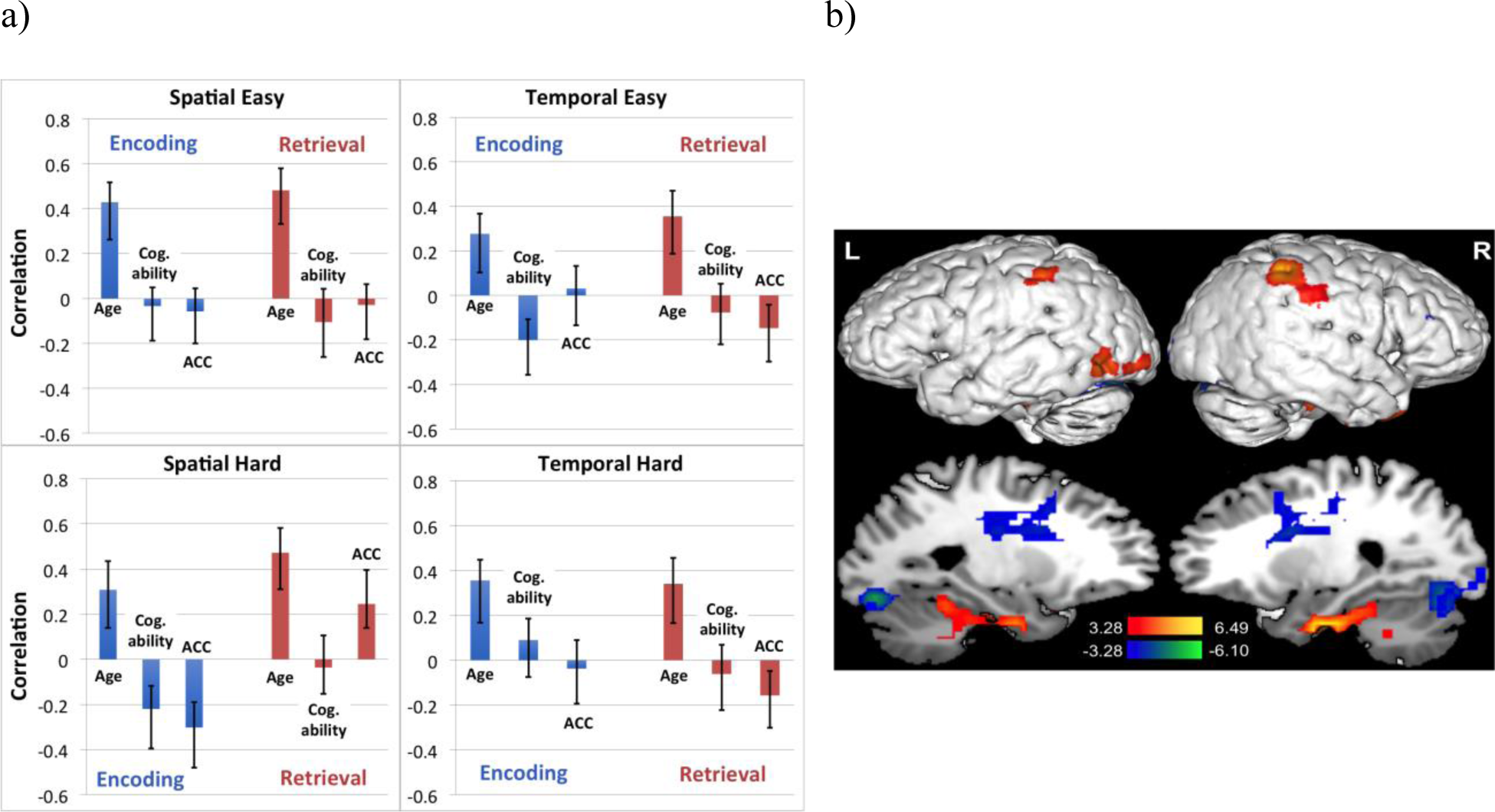
Brain-behaviour correlation profile and corresponding singular image for LV2 in cognitive ability B-PLS. a) LV2 brain-behaviour correlation profile for cognitive ability B-PLS separated by task. The correlation profile indicated that activity in positive salience regions increased with age, and activity in negative salience regions decreased with age across tasks. ACC is short for accuracy, and Cog. ability is short for cognitive ability. Error bars represent 95% confidence intervals. b) Singular image for LV2 of cognitive ability B-PLS showing positive (warm coloured regions) and negative (cool coloured regions) voxel saliences. The scale represents the range of bootstrap ratio values thresholded at ± 3.28, p < 0.001. Activations are presented on template images of the lateral and medial surfaces of the left and right hemispheres of the brain using Multi-image Analysis GUI (Mango) software (http://ric.uthscsa.edu/mango/). The effects in LV2 of cognitive ability B-PLS replicated the same pattern of age-related increases and decreases of activity with age across all encoding and retrieval trials discussed previously in LV2 of the reserve B-PLS analysis. The current LV accounted for 12.27% of the total cross-block covariance (p < 0.001). The post-hoc GLM for brain scores against age, cognitive ability, and task accuracy (R2 = 0.14, F (3,1228) = 69.38, p < .001) revealed a significant main effect for age across task conditions (p < 0.001). The full list of brain regions from this LV is outlined in Supplementary Table 1.

**Supplementary Figure 2.**
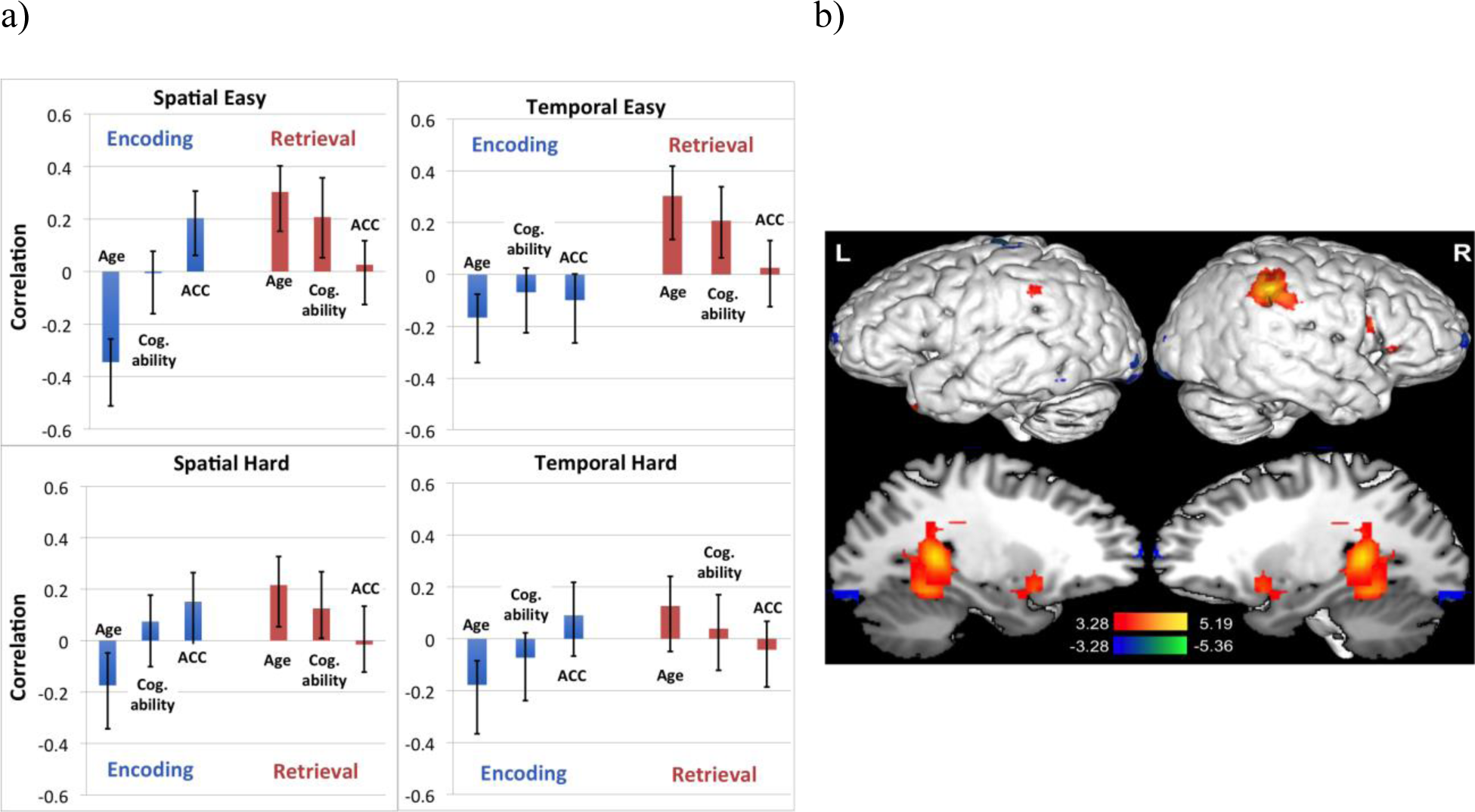
Brain-behaviour correlation profile and corresponding singular image for LV3 of cognitive ability B-PLS analysis. a) LV3 brain-behaviour correlation profile for cognitive ability B-PLS separated by task. The correlation profile indicated that activity in positive salience regions increased with age at retrieval and decreased with age at encoding. Activity in negative salience regions increased with age at encoding and decreased with age at retrieval. Activity in negative salience brain regions also increased with cognitive ability (Cog. ability) at retrieval (except for TH). ACC is short for accuracy. Error bars represent 95% confidence intervals. b) Singular image for LV3 of cognitive ability B-PLS showing positive voxel saliences (warm coloured regions) and negative voxel saliences (cool coloured regions). The scale represents the range of bootstrap ratio values thresholded at ± 3.28, p < 0.001. Activations are presented on template images of the lateral and medial surfaces of the left and right hemispheres of the brain using Multi-image Analysis GUI (Mango) software (http://ric.uthscsa.edu/mango/). The pattern revealed in LV3 of the cognitive ability B-PLS mirrored the age × phase interaction discussed in LV3 of the reserve B-PLS (i.e., brain regions that were differentially related to age during encoding and retrieval). This LV accounted for 10.15% of the total cross-block covariance. In this analysis, in addition to the positive association between age and activity in positive salience brain during retrieval, activity in those regions was also positively correlated with cognitive ability (except during TH). Negative salience regions were also negatively correlated with cognitive ability, during retrieval (except for TH). Local maxima of the positive and negative salience brain regions in this LV are fairly similar to the ones observed in LV3 of the first analysis. Post-hoc GLMs for brain scores against age, cognitive ability, and task accuracy at encoding (R^2^ = 0.05, F (3, 612) = 11.29, p < .001), and retrieval (R^2^ = 0.07, F (3, 612) = 15.44, p < .001), revealed significant main effects of age (ps < 0.01), but in opposite directions, mirroring the age × phase interaction outlined in the PLS correlation profile. Additionally, post hoc GLM revealed a significant main effect of cognitive ability, but only at retrieval (p < 0.01). The full list of brain regions from this LV is outlined in Supplementary Table 2.

**Supplementary Table 1:**
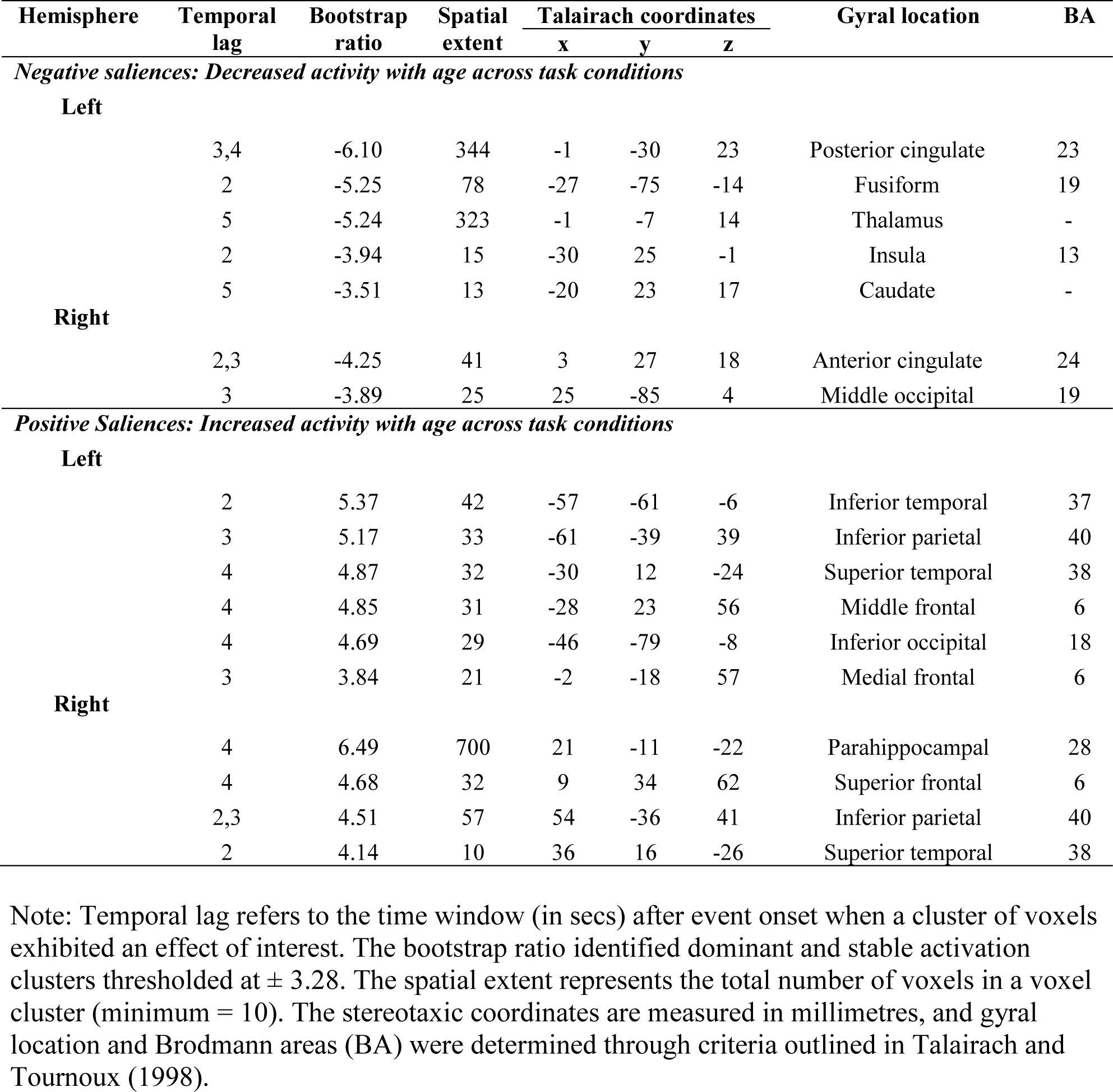
Local maxima for LV2 of cognitive ability B-PLS: Regions where activity correlated with age across encoding and retrieval phases

**Supplementary Table 2:** Local maxima for LV3 of cognitive ability B-PLS: regions where activity correlated with age differentially at encoding and retrieval

| Hemisphere | Temporal lag | Bootstrap ratio | Spatial extent | Talairach coordinates |  |  | Gyral location | BA |
| --- | --- | --- | --- | --- | --- | --- | --- | --- |
|  |  |  |  | x | y | z |  |  |
| *Negative Saliences: Increased activity with age at encoding and decreased at retrieval, decreased with cognitive ability during retrieval (except for temporal hard)* |  |  |  |  |  |  |  |  |
| Left |  |  |  |  |  |  |  |  |
|  | 2,3,5 | -5.36 | 79 | -35 | -34 | 62 | Postcentral | 1,2,3 |
|  | 2,3,5 | -4.78 | 27 | -13 | -68 | 30 | Precuneus | 7 |
|  | 3 | -4.49 | 14 | -23 | 57 | 20 | Middle frontal | 10 |
|  | 5 | -4.34 | 17 | -57 | -64 | -10 | Middle/inferior occipital | 18/37 |
|  | 3 | -4.08 | 35 | -27 | -75 | -14 | Fusiform | 19 |
| Right |  |  |  |  |  |  |  |  |
|  | 2 | -4.70 | 19 | 29 | 57 | 17 | Superior frontal | 10 |
|  | 3,4,5 | -4.66 | 38 | 25 | -90 | -15 | Fusiform/lingual | 18/37 |
|  | 5 | -4.57 | 49 | 46 | -7 | 55 | Precentral | 6 |
|  | 5 | -3.64 | 11 | 51 | -15 | -21 | Middle temporal | 21 |
| *Positive Saliences: Increased activity with age at retrieval and decreased at encoding, and increased with cognitive ability during retrieval (except for temporal hard)* |  |  |  |  |  |  |  |  |
| Left |  |  |  |  |  |  |  |  |
|  | 2 | 5.19 | 10 | -38 | 20 | -27 | Superior temporal | 38 |
|  | 5 | 5.07 | 166 | -27 | -43 | 7 | Hippocampus | - |
|  | 2 | 4.10 | 25 | -23 | 7 | -6 | Putamen | - |
|  | 3 | 3.94 | 17 | -50 | -42 | 28 | Inferior parietal | 40 |
| Right |  |  |  |  |  |  |  |  |
|  | 5 | 5.08 | 187 | 29 | -39 | 1 | Hippocampus/<br>Parahippocampus | - |
|  | 2 | 4.61 | 31 | 21 | 10 | -2 | Putamen | - |
|  | 3 | 4.18 | 19 | 51 | 4 | 20 | Inferior frontal | 44 |
|  | 5 | 3.92 | 23 | 36 | -46 | 29 | Inferior parietal | 40 |

Note: Temporal lag refers to the time window (in secs) after event onset when a cluster of voxels exhibited an effect of interest. The bootstrap ratio identified dominant and stable activation clusters thresholded at $\pm 3.28$ . The spatial extent represents the total number of voxels in a voxel cluster (minimum = 10). The stereotaxic coordinates are measured in millimetres, and gyrus location and Brodmann areas (BA) were determined through criteria outlined in Talairach and Tournoux (1998).

## Notes

#### Summary of Updates

We re-ran our B-PLS analysis using (i) reserve and (ii) cognitive ability as vectors of interest in two separate analyses. We then compared similarities and differences in the two analyses.

